# Structural Basis for the Improved RNA-Clamping of Amidino-Rocaglates to eIF4A1

**DOI:** 10.1101/2024.10.04.616674

**Authors:** James F. Conley, Lauren E. Brown, James H. McNeely, Jerry Pelletier, John A. Porco, Karen N. Allen

## Abstract

Eukaryotic Initiation Factor 4A-1 (eIF4A1) is an ATP-dependent RNA helicase that unwinds 5’-UTR mRNA secondary structures to facilitate cap-dependent translation initiation. Rocaglates, a class of natural products typified by rocaglamide A (RocA), possess anti-neoplastic and anti-infectious activity mediated by their interaction with eIF4A1. Rocaglates inhibit cap-dependent translation initiation by “clamping” eIF4A1 onto polypurine RNA, impeding ribosome scanning. A novel class of rocaglate derivatives, amidino-rocaglates (ADRs) which feature an amidine ring fused to the rocaglate core, are particularly effective at promoting eIF4A1-RNA-clamping compared to other rocaglate congeners. Herein, we present the X-ray crystal structure of an ADR in complex with eIF4A1, the nonhydrolyzable ATP ground-state mimic adenylyl-imidodiphosphate (AMPPNP), and poly r(AG)_5_ RNA refined to 1.69 Å resolution. The binding pose and interactions of the ADR with eIF4A1 do not differ substantially from those of RocA, prompting investigation of the basis for enhanced target engagement. Computational modeling suggests that the rigidified ADR scaffold is inherently preorganized in an eIF4A1-RNA-binding-competent conformation, thereby avoiding entropic penalties associated with RocA binding. This study illustrates how conformational rigidification of the rocaglate scaffold can be leveraged to improve potency for the development of rocaglates as potential anticancer and anti-infectious agents.

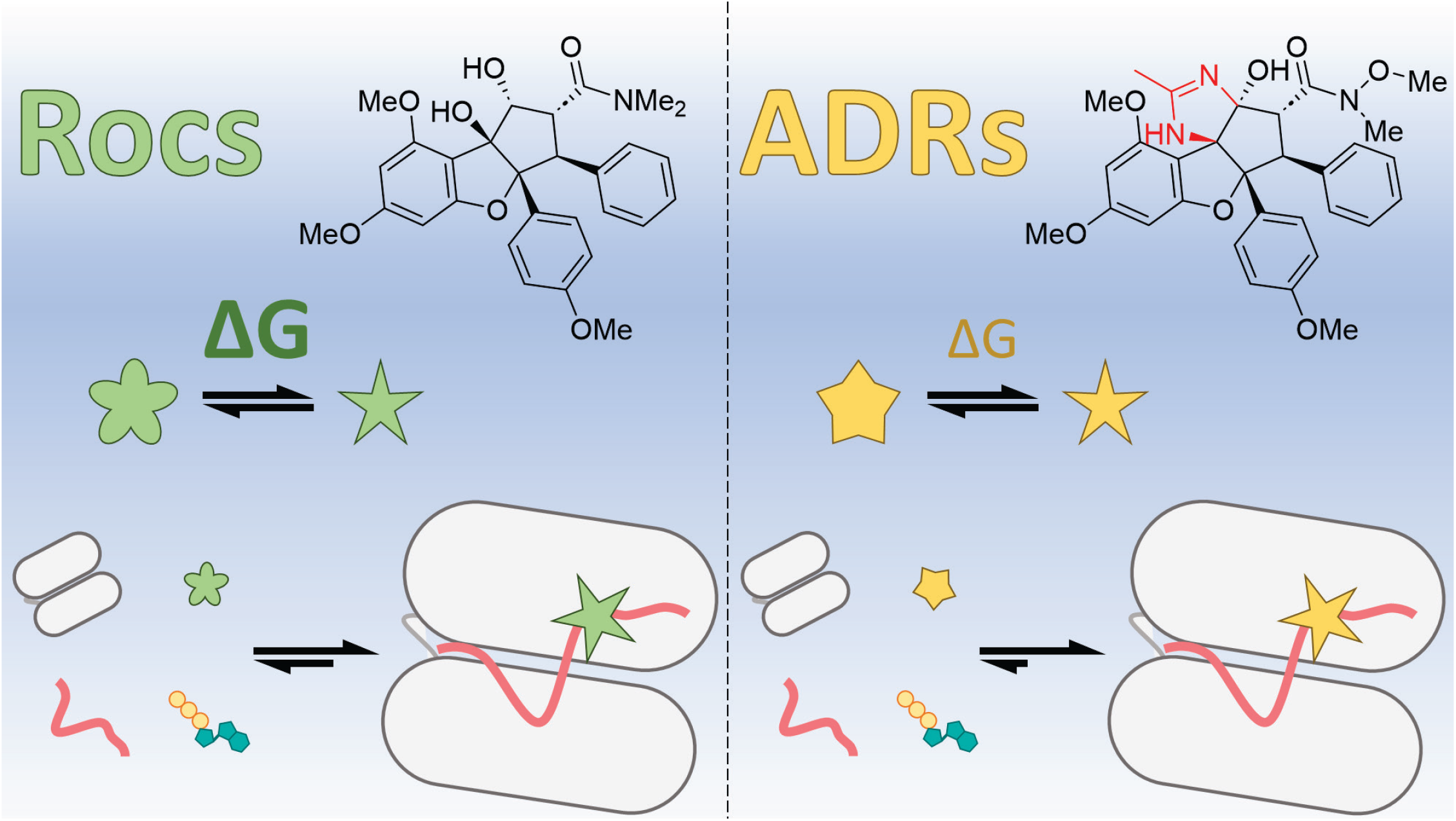

## Introduction

Protein translation is arguably the most critical cellular process and is extensively regulated, requiring the involvement of dozens of proteins to modulate protein synthesis both temporally and spatially. Translation is comprised of three sequential steps: initiation, elongation, and termination. Initiation involves the recruitment of the cellular translation machinery to mRNA and is the primary point of regulation within the translation process.^1^ The majority of eukaryotic initiation proceeds though a “cap-dependent” mechanism, denoted for the importance of the 5’-7-methylguanosine (m^7^G) cap on the nascent mRNA strand for recognition by <u>e</u>ukaryotic <u>i</u>nitiation <u>f</u>actors (eIFs). Different mRNAs possess varying degrees of secondary structure that require the recruitment of eIFs to unwind the RNA and facilitate efficient ribosome scanning for the initiation codon by the 43S preinitiation complex. eIF4A is the RNA helicase responsible for unwinding secondary structure in the 5’-untranslated regions (UTRs) of mRNAs and is a key component of the eIF4F cap-binding complex along with the 5’-m^7^G cap-binding protein eIF4E and the scaffolding protein eIF4G.

In humans and most higher eukaryotes, there are three paralogs within the eIF4A subfamily of DEAD box helicases: eIF4A1, eIF4A2, and eIF4A3. eIF4A1 is the predominant paralog involved in translation initiation, whereas eIF4A2 and eIF4A3 are involved in other processes related to mRNA metabolism.^2^ eIF4A1 (DDX2A) and eIF4A2 (DDX2B) are very similar (90% sequence identity), and though they have been shown to be functionally interchangeable *in vitro*, the two proteins have distinct functions *in vivo* and are expressed at varying levels in different cell types.^3,4^ Furthermore, overexpression of either eIF4A1 or eIF4A2 cannot compensate for the loss of the other for their respective functions.^5^ eIF4A3 (DDX48) is more divergent and is functionally distinct from the other eIF4A paralogs.^2^ eIF4A1 is one of the most ubiquitously expressed enzymes in eukaryotic cells and is a crucial node of the initiation step of eukaryotic cap-dependent translation.^6,7^ Modulation of RNA unwinding through the direct regulation of eIF4A1 activity is a critical post-transcriptional means of not only modulating the rate of global protein translation, but also regulating which mRNA transcripts are translated. eIF4A1 overexpression has been associated with poor outcomes in several cancers and, as a result, much of the recent research on it is contextualized by the investigation of its role in cancer and as a target for inhibition.^8^

Pharmacologic inhibition of eIF4A by several natural products, with varying degrees of specificity for the three paralogs, has been shown to have a potent anti-neoplastic effect in both *in vitro* and *in vivo* models of cancer.^5,8–12^ Rocaglates are one example of a natural product class with a unique mechanism of selective inhibition of the translation of purine-rich mRNA mediated by their interaction with eIF4A1.^13–15^ Rocaglates are a class of natural products of phytochemical origin that contain a cyclopenta[b]benzofuran core. The prototypical rocaglate, rocaglamide A (RocA), was first identified and isolated in 1982 from extracts of plant *Aglaia elliptifolia*.^16^ Since then, hundreds of related compounds, including silvesterol, have been isolated from various members of the genus *Aglaia*, and efficient syntheses have been developed to expand this chemical space (**Supplementary Figure S1**).^17,18^ Zotatifin (eFT226) is one such synthetic rocaglate shown to be an effective inhibitor of tumor growth in receptor tyrosine kinase-driven cancers that is currently being evaluated in the clinic as a component of combination therapies against estrogen receptor positive metastatic breast cancer.^19,20^

Iwasaki and coworkers have published a pair of articles characterizing the mechanism of action of RocA, both biochemically and structurally.^15,16,21^ RocA was shown to diminish protein production in a variety of model systems in a dose-dependent manner, with a notable difference in the magnitude of translational repression of 5’-UTRs containing high purine content. Such transcripts were found to be much more susceptible to translation repression by RocA than transcripts with 5’-UTRs containing elevated pyrimidine content.^15^ The X-ray crystal structure of RocA bound to eIF4A1 and purine RNA revealed the rocaglate binding site at the interface of eIF4A1 and polypurine RNA, demonstrating how rocaglates force engagement between protein and RNA and lock the helicase into a closed conformation in a mechanism termed “clamping.”^21^ In this clamped state, eIF4A1 can no longer undergo the multiple rounds of RNA remodeling required for efficient start codon scanning, eIF4A1 is depleted from eIF4F, translation initiation is inhibited, and protein production in the cell is diminished.^15,21^ The sequence selectivity of rocaglates is thought to arise from π-stacking interactions between the rings A and B on the rocaglate core and consecutive purine nucleobases, along with a critical hydrogen bond between 8b-OH of RocA and N7 of a purine nucleobase, guanine 8 (G8) in the case of the published structure (**Figure 1A** and **Figure 2**).^21^ When polypyrimidine RNA is modeled in place of polypurine RNA, the relevant components of π-stacking interactions are misaligned and there is no hydrogen-bond acceptor for 8b-OH at the same location on the pyrimidine nucleobase.^21^ This important structure showed additional details of target engagement, including a π-stacking interaction between the C ring of RocA and F163 and a hydrogen bond between the carbonyl of RocA’s *C*2-*N,N*-dimethyl-carboxamide and the side-chain NH_2_ group of Q195.^21^

**Figure 1.**
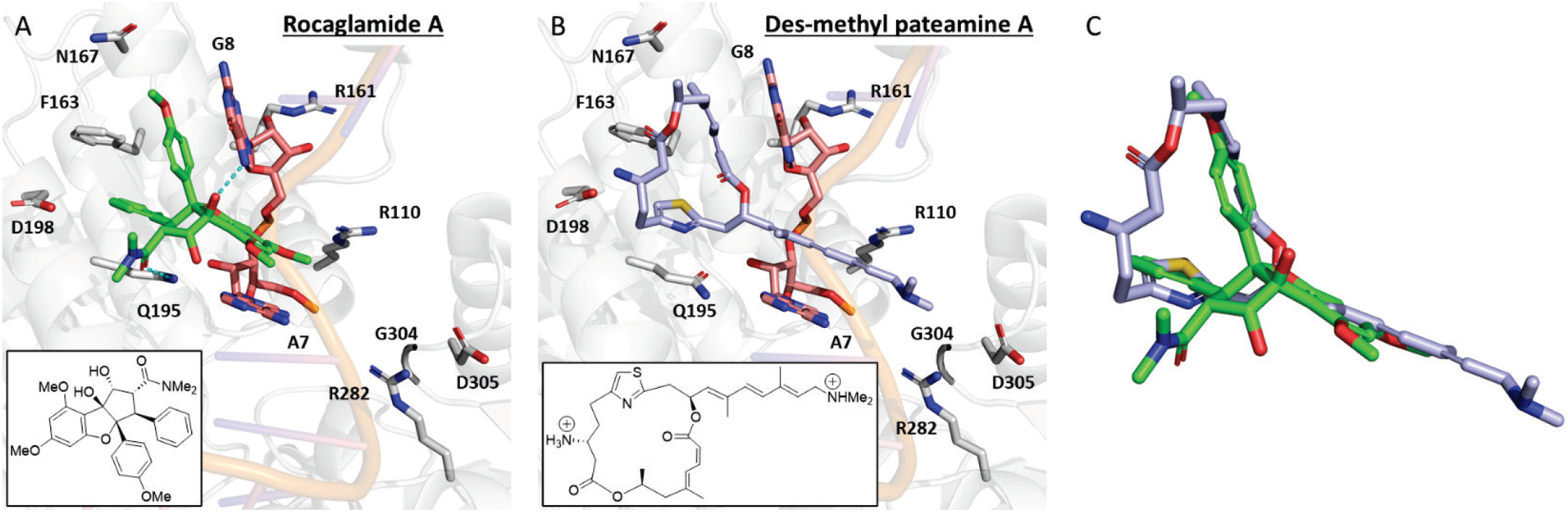
Comparison of the structures of eIF4A1:inhibitor complexes. **A**. eIF4A1:RocA:RNA:AMPPNP complex from 5ZC9 with corresponding chemical structure of RocA (inset). **B**. eIF4A1:RocA:RNA:AMPPNP complex from 6XKI with corresponding structure of des-methyl pateamine A (DMPatA; MZ-735, inset). Important residues for inhibitor binding are labeled. **C**. RocA overlaid with DMPatA by superposition of liganded structures.

**Figure 2.**
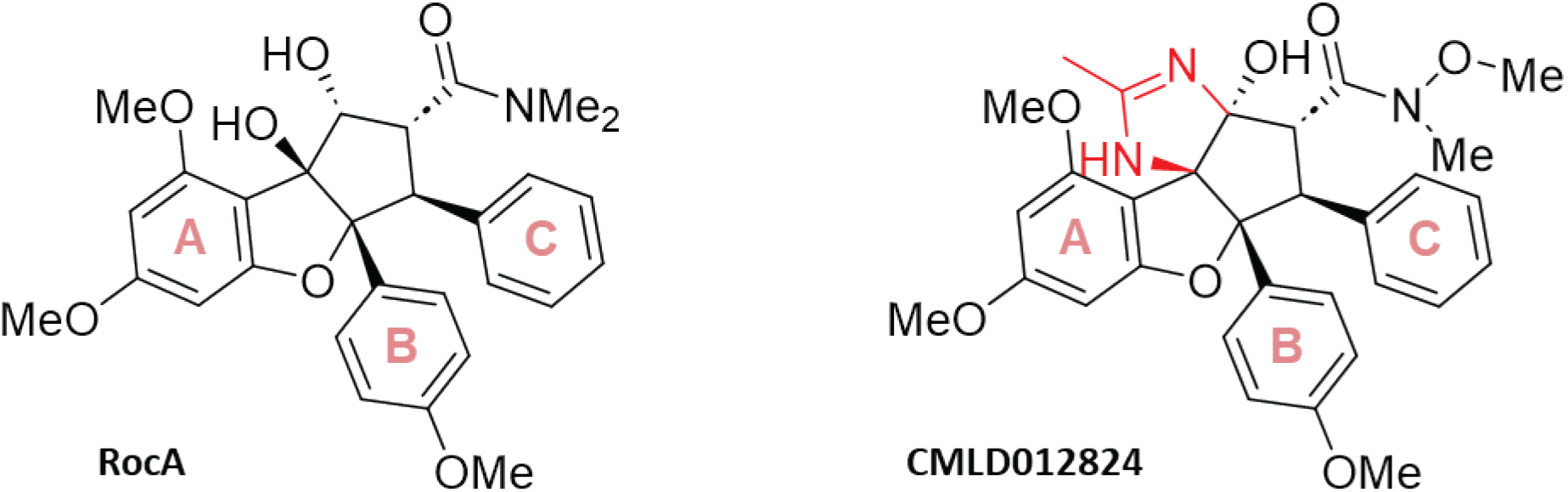
Chemical structure of RocA and ADR CMLD012824. RocA, the first rocaglate described, and **CMLD012824**, a potent derivative from a new class of compounds – amidino-rocaglates (ADRs)– share the same cyclopenta[b]benzofuran core with the A, B, and C rings labeled on each structure. ADRs feature an annulated heterocyclic amidine ring (shown in red) fused to the rocaglate scaffold.

A recent paper by the Pelletier and Romo groups reported the X-ray crystal structure of eIF4A1 bound to another translation inhibitor of a different chemical class, desmethyl pateamine A (DMPatA), a derivative of the marine sponge-derived natural product pateamine A (PatA).^22^ Pateamines differ significantly from rocaglates in both chemical structure and pharmacophoric elements; these compounds are macrodiolides comprised of a decorated 19-membered ring fused to a thiazole ring, with an additional large conjugated alkyl chain appended in a lariat-type structure. DMPatA differs from PatA by the removal of a single methyl group on the macrocycle. Despite a lack of structural similarity, DMPatA and RocA bind at the same location at the eIF4A1-RNA interface (**Figure 1B**).^22^ The macrocycle of DMPatA occupies the same pocket as the B and C rings of RocA, while the conjugated alkyl chain occupies the same space as the A ring, while protruding further towards residues R282, G304, and D305 on the surface of eIF4A1 (**Figure 1B** and **Figure 1C**). Notably, DMPatA does not have a unique interaction with purine nucleobases, explaining its superior capacity to inhibit translation of pyrimidine-rich mRNAs compared to RocA. Nevertheless, it does exhibit increased clamping activity for purine-rich mRNAs, similar to RocA; this difference can possibly be explained by the increased buried surface area arising from binding in complex with purine nucleotides.^22^ As the authors describe in their title, DMPatA seems to exhibit a unique “functional mimicry” to rocaglates, suggesting that the stabilization of a preferred clamped orientation of RNA bound to eIF4A1 is of greater importance than the specific chemical structure of eIF4A-clamping inhibitors, like rocaglates or PatA analogs. These studies together suggest that diverse pharmacophores can be deployed to clamp RNA to eIF4A1 and inhibit downstream protein synthesis, and that broad chemical space is available for generation of helicase-interacting compounds that operate by a “clamping”-type mechanism.

We have previously reported amidino rocaglates (ADRs), a class of synthetic rocaglates with an amidine ring fused to the cyclopenta[*b*]benzofuran scaffold, which were serendipitously obtained from a retro-Nazarov reaction performed on a tosylated rocaglate core.^23^ In our collaborative studies, ADRs were found to have exceptionally strong eIF4A clamping to polypurine RNA and exhibit potent cap-dependent translation inhibition.^24,25^ Moreover, ADRs and a related guanidino rocaglate (GDR) were found to possess strong antiviral activity against hepatitis E virus (HEV) with EC_50_ values between 1 and 9 nM.^26^ The strong potency of ADRs and emerging biology led us to undertake structural biology studies to probe the nature of heightened target engagement.

Herein, we report the X-ray crystal structure, refined to 1.69 Å resolution, of an exemplar ADR, the potent translation inhibitor **CMLD012824**, in complex with eIF4A1, the nonhydrolyzable ATP analog adenylyl-imidodiphosphate (AMPPNP), and a poly r(AG)_5_. This structural information combined with computational analysis suggests that conformational preorganization provided by the ADR-ring fusion is a key contributor to the enhanced activity on the rocaglate scaffold, suggesting new avenues for the development of inhibitory DEAD Box helicase ligands.

## Results and Discussion

### Structure determination of an ADR-eIF4A-RNA-AMPPNP co-crystal complex

The main purpose of our investigation was to characterize the binding mode of ADRs and identify potential molecular determinants of their improved inhibition and target engagement of the eIF4A1-polypurine RNA interface. The ADR chosen for study was **CMLD012824**, an enantioenriched preparation of racemic **CMLD012612**, that is the most potent ADR identified to date.^24^ To this end, we sought to determine the structure of the assumed quaternary complex formed by eIF4A1 bound to polypurine RNA, an ATP analog, and the ADR. Each component of the quaternary complex is critical for the formation of a stable homogeneous crystallizable unit in which eIF4A1 adopts and maintains a closed conformation. The previously reported structure of the RocA-bound quaternary complex was used as a reference for protein preparation. In that structure, HRV-3C cleaved eIF4A1(19-406) was used for crystallization experiments, however only residues 29-406 are well-resolved, offering the potential for more liberal truncation of the N-terminus. Based on this observation, we designed a smaller crystallization construct with a cleavable N-terminal SUMO-tag and a short -Ser-Ser-linker followed by residues 21-406. Rocaglates require the association of purine RNA to eIF4A1 for binding.^21^ The polypurine RNA selected was poly r(AG)_5_. eIF4A1, like other DEAD-box helicases, requires Mg^2+^-ATP binding for RNA binding and ATP hydrolysis for successive unwinding events; thus, a non-hydrolysable analog, AMPPNP, was included.^6,27,28^ Poly r(AG)_5_ and AMPPNP were each used in the published structure of RocA, so represented a reasonable starting point for this effort.

eIF4A1 crystallized in complex with **CMLD012824**, AMPPNP, and poly r(AG)_5_ in the space group P1 with four of each molecule in the asymmetric unit (ASU), which in this space group is equivalent to the unit cell (**Figure 3; Table 1**). This structure was refined to 1.69 Å (R_work_/R_free_: 0.20/0.23). The ASU exhibits pseudo-twofold rotational symmetry about its center, with two complexes modeled with their protein-RNA interfaces facing toward each other, and each of the other two complexes rotated ∼90° to their corresponding pseudo-symmetry mate positioned at the interface of the two central protein chains (**Supplementary Figure S2**). A notable difference between this structure and the previously published RocA complex is the manner in which the RNA oligonucleotide binds. In the crystal structure of the RocA complex, each nucleotide in the 10-mer oligonucleotide has electron density which is well-resolved, whereas in this structure, only 8 of the 10 nucleotides in each chain are well-resolved. When examining the crystal contacts and solvent channels in the **CMLD012824** structure, there is not sufficient space to build two additional nucleotides at the 5’ end of the RNA strands. This suggests that RNA binding in the **CMLD012824** complex is shifted by two nucleotide positions in order to allow packing into this crystal form. Thus, what was previously described as π-stacking between A7 and G8 of the oligonucleotide with Ring A and Ring B of RocA in that complex, is assigned as π-stacking between A5 and G6 of the oligonucleotide with Ring A and Ring B of **CMLD012824** (**Figure 4**). The most notable difference between the pairs of pseudo-symmetry mates is the ring-stacking interactions of the 5’ end of the RNA strands. Whereas in Chains X and Z the A1 nucleobases from each strand interact, in chains W and Y the A1 nucleobases stack with the A2 nucleobases of the other RNA strand (**Supplementary Figure S3**). Additionally, the loop containing Motif VI common to DEAD-box helicases (G363-G370) that directly interacts with the AMPPNP ligand is apparently positioned differently across some of the chains.^6^ In Chains A and C, these loops adopt very similar conformations to one another, consistent with the canonical ATP-bound conformation, while in Chain B, density for the loop is clearly resolved in an alternate conformation, notably for ATP-interacting residues R365 and F366, and in chain D the entire loop is poorly resolved (data not shown). We do not predict these differences to have any impact on **CMLD012824** binding, as each AMPPNP ligand is well resolved in nearly identical position when the chains are aligned, and thus the quaternary complex appears unaffected. Other minor differences between the chains include the extent to which the N- and C-terminal residues can be resolved, namely there is clear electron density for Chain A 29-406, Chain B 26-400, Chain C 29-406 and Chain D 26-399. Otherwise, each quaternary complex overlays with the others with low root-mean square deviation (RMSD) of 0.079-0.197 Å^2^.

**Table 1.** Data collection and refinement statistics.

|  | Human eIF4A1/AMPPNP/CMLD012824/poly(AG)<sub>5</sub> complex<br>PDB: 9DTS |
| --- | --- |
| Beamline | NSLS-II 17-ID-1 (AMX), BNL |
| Wavelength | 0.9201 |
| Resolution range | 36.51 - 1.69 (1.75 - 1.69) |
| Space group | P 1 |
| Unit cell (a, b, c)<br>($\alpha$, $\beta$, $\gamma$) | 66.11, 87.314, 93.159<br>95.222, 105.287, 108.353 |
| Total reflections | 597134 (52543) |
| Unique reflections | 201775 (19049) |
| Multiplicity | 3.0 (2.7) |
| Completeness (%) | 96.00 (91.16) |
| Mean I/sigma(I) | 5.59 (0.80) |
| Wilson B-factor | 27.27 |
| R-merge | 0.09026 (1.177) |
| R-meas | 0.1095 (1.454) |
| R-pim | 0.06117 (0.8404) |
| CC1/2 | 0.993 (0.368) |
| CC* | 0.998 (0.733) |
| Reflections used in refinement | 201334 (19047) |
| R-work | 0.2028 (0.3432) |
| R-free | 0.2400 (0.3363) |
| CC(work) | 0.943 (0.638) |
| CC(free) | 0.924 (0.575) |
| Number of non-hydrogen atoms | 14030 |
| macromolecules | 12839 |
| ligands | 472 |
| solvent | 899 |
| Protein residues | 1503 |
| RMS(bonds) | 0.012 |
| RMS(angles) | 1.25 |
| Ramachandran favored (%) | 97.73 |
| Ramachandran allowed (%) | 1.87 |
| Ramachandran outliers (%) | 0.40 |
| Rotamer outliers (%) | 1.27 |
| Clashscore | 8.58 |
| Average B-factor | 40.02 |
| macromolecules | 39.95 |
| ligands | 28.70 |
| solvent | 41.81 |

Statistics for the highest-resolution shell are shown in parentheses.

**Figure 3.**
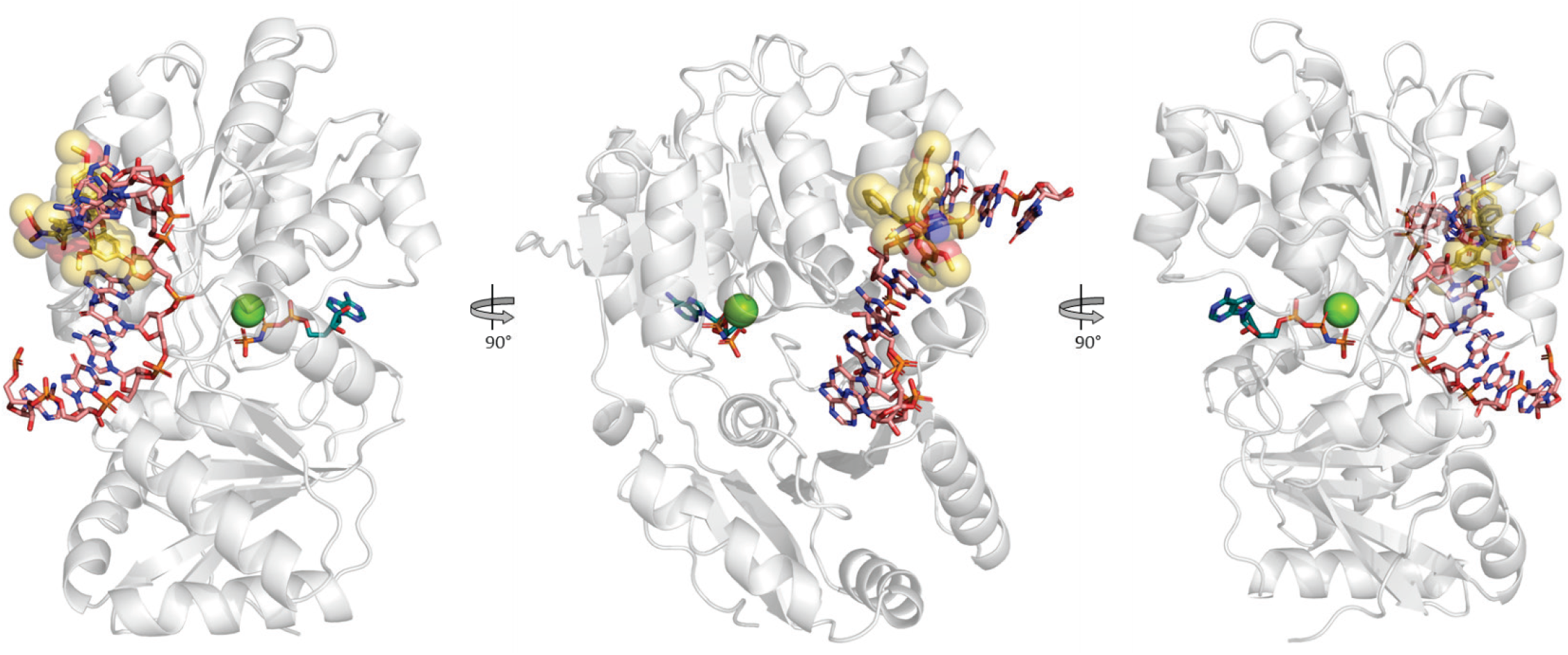
Structure of the eIF4A1:CMLD012824:RNA:AMPPNP quaternary complex. eIF4A1 shown as off-white transparent ribbon with poly (AG)_5_ (salmon) **CMLD012824** (yellow stick with transparent spheres), AMPPNP (teal) shown as sticks and Mg^2+^ as green sphere.

**Figure 4.**
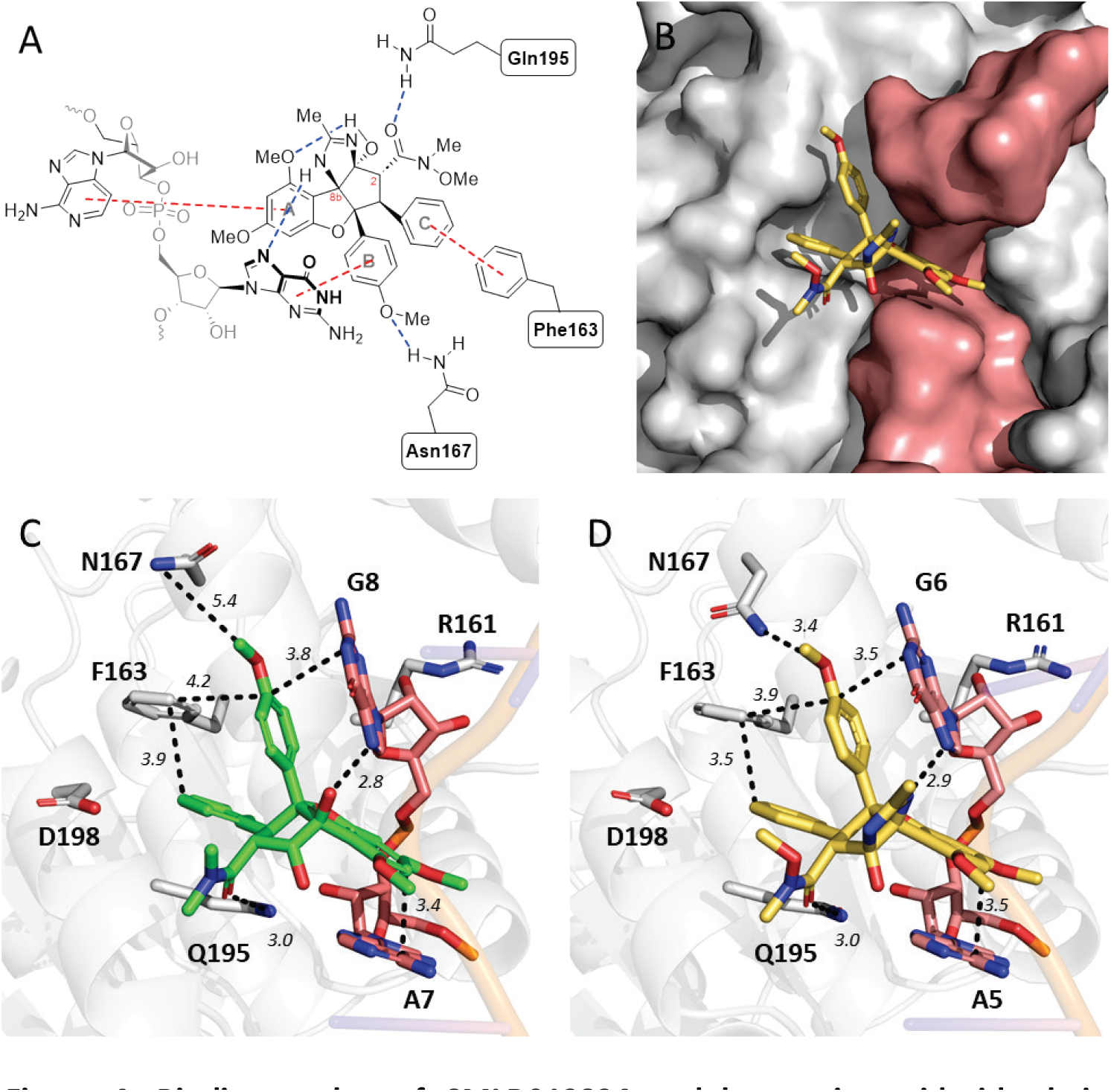
Binding pocket of CMLD012824 and key amino-acid side-chain interactions. **(A)** Interaction diagram for bound eIF4A1-poly (AG)_5_-CMLD012824. **(B)** Surface representation of interface of eIF4A1 (white) and RNA (salmon) comprising binding pocket for CMLD012824 (yellow stick). Key interactions for RocA **(C)** and CMLD012824 **(D)**, with hydrogen bond and hydrophobic interactions as dashed lines with lengths in Å. eIF4A1, off-white; poly (AG)_5_, salmon; RocA, green; CMLD012824, yellow.

The positioning and binding mode of **CMLD012824** is nearly identical to that of RocA from the previously reported structure, and consistent with the predicted binding mode from our previously published modeling.^21,24^ The binding pocket for **CMLD012824** is formed at the protein-RNA interface, with the ADR positioned between consecutive purine nucleobases of the kinked RNA strand and with the C-Ring projecting into a pocket formed at the interface of two α-helices (**Figure 4**). The presence of polypurine RNA is essential to the formation of the binding pocket. FTMap, a computational mapping server that identifies binding “hot spots,” or surfaces with a high propensity for ligand binding, did not identify the rocaglate binding site as a hot spot in the absence of the RNA, despite identifying the ATP-binding site and portions of the RNA binding sites.^29^ As in the case of RocA, aryl rings A, B, and C (**Figure 1**) stack with A5, G6, and the sidechain phenyl ring of F163, respectively, at distances between 3.6-4.0 Å (**Figure 4**). Unique to ADRs, the imidazoline N-H is engaged in a hydrogen bond to *N7* of G6. The distance and positioning of this hydrogen bond is comparable to that observed between C8b tertiary hydroxyl of RocA and *N7* of G8 (2.7-2.9 Å, depending on the chain), which is thought to contribute to the unique purine selectivity of rocaglate binding.^24^ In the present structure, the B ring methoxy group and N167 are within hydrogen bonding distance, which is not the case in the previously reported structure of RocA, due to a conformational difference in the N167 sidechain (**Figure 4**). This first crystallographic observation of the formation of a hydrogen bond at this position and reaffirms the importance of a hydrogen bond acceptor at the *para*-position of the B ring of rocaglates for the potency of these compounds.^30^ Hydrogen-bonding interactions provide a decrease in free energy that is offset by desolvation penalties; the high degree of solvent exposure at this site suggests that any gains to binding affinity imparted by this newly observed H-bond would be modest at best. Thus, we do not expect this to fully explain the improved RNA-clamping activity observed for **CMLD012824**. Moreover, the position of the B ring does not differ between RocA and **CMLD012824**, so it is reasonable to assume that this hydrogen bond could also be made between eIF4A1 and RocA with a slight conformational adjustment of the protein. Given the considerable similarity between the binding interactions observed between the protein and each of the compounds, we next sought to use computational modeling to rationalize the substantial gains in RNA-clamping and translation inhibition activity of ADRs over traditional rocaglates.

### Computational modeling of rocaglate ligands

In prior studies, we briefly described efforts to model **CMLD012824** into the published RocA X-ray structure using a manual-overlay.^24^ To leverage our new structural data and improve upon this earlier modeling, we performed computational docking (Glide, Schrodinger LLC) of **CMLD012824** and RocA into their respective X-ray structures.^31,32^ The purpose of this docking was to determine whether the docking scores, which serve to approximate the predicted free energy of binding (ΔG_binding_), would reveal any meaningful differences between the **CMLD012824** and RocA ligands. For both compounds, the top computationally predicted poses were in good agreement with the experimentally-determined binding modes (**Figure 5**). In the case of **CMLD012824**, however, the *N,O*-dimethylhydroxylamine moiety of the *C*2-tethered Weinreb amide was predicted to be flipped approximately 180° relative to the experimentally-determined pose (**Figure 5B**); given the lack of protein or RNA interactions and solvent exposure of these atoms we consider this difference to have a negligible impact on binding energy. The Glide Gscores for **CMLD012824** and RocA docked into their native X-ray structures were quite similar (−11.1 vs -11.8 kcal/mol, respectively), and while the score for RocA was slightly improved relative to **CMLD012824**, limitations in the predictive accuracy of Glide docking (RMSD 2.3 kcal/mol) precludes a meaningful head-to-head comparison of these energies.^31^ As such, these docking results support our empirical observations that RocA and **CMLD012824** each engage eIF4A/RNA in a highly similar manner, with analogous binding interactions that are not expected to lead to substantial enthalpic differences in binding affinity.

**Figure 5.**
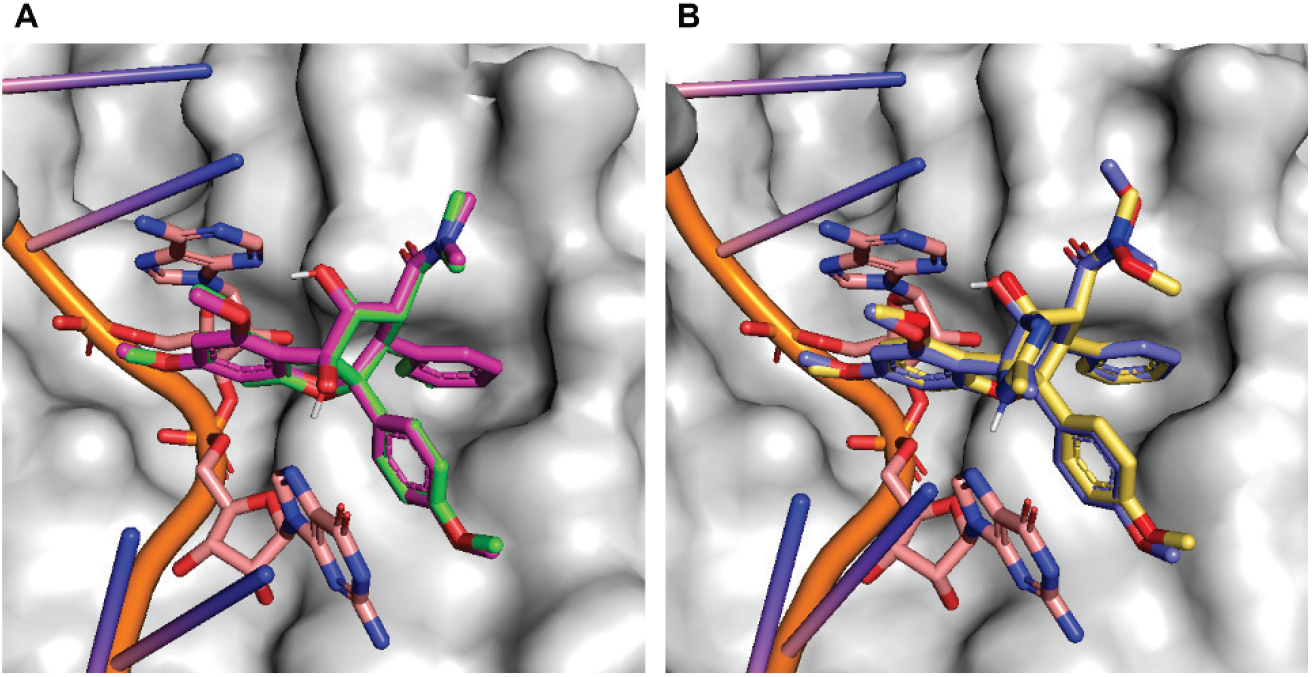
Computational docking of RocA and CMLD012824 show high agreement with the experimentally determined structures, without substantial differences in predicted binding energies. **(A)** Glide-predicted docking pose (G_score_: -11.8 kcal/mol) for RocA (*magenta*) docked into the 5ZC9 eIF4A:RNA structure (*white*/*salmon*) and overlaid with liganded RocA (*green*) from this structure. **(B)** Glide-predicted docking pose (G_score_: -11.1 kcal/mol) for **CMLD012824** (*lavender*) docked into the eIF4A1-poly (AG)_5_ structure (*white*/*salmon*) and overlaid with liganded **CMLD012824** from this structure (*gold*).

Given these observations, we hypothesized that the conformational constraints imposed by the fused amidine ring may contribute to improved target engagement via conformational preorganization in a “binding-competent” mode, leading to reduced entropic penalties on binding. Conformational restriction by way of bond desaturation, ring formation, or other rigidification strategies such as intramolecular hydrogen bonding is a commonly deployed medicinal chemistry and structure-based drug design strategy to enhance compound potency by way of reducing the entropic term (TΔS) which offsets enthalpic gains (ΔH) achieved by ligand-target interactions.^33–40^ We noted that the **CMLD012824** amidine N-H, which from our X-ray crystal structure is proposed to mimic the RocA C8b hydroxyl in hydrogen bonding an RNA guanine base (G8/G6, respectively), is conformationally locked in the orientation required for hydrogen-bond donation. This contrasts the untethered RocA hydroxyl, which can freely rotate to allow for competing hydrogen bonds with solvent (water) in the unbound state. This observation led to the question of whether the rigidifying effect of the amidine fusion on the already-constrained cyclopenta[*b*]benzofuran rocaglate core may also impact the positioning of other key pendant functional groups with respect to their protein/RNA binding partners – namely the Q195-interacting C2 substituent, the adenine-interacting A-ring, the guanine-interacting B-ring, and the Phe163 interacting C-ring. For example, during the development of rocaglate clinical candidate eFT226 (Zotatifin), Ernst and coworkers utilized DFT modeling of RocA to identify an optimal ∼40° torsion between the pendant “B” and “C” rocaglate rings that was deployed in synthetic candidate selection and was later recapitulated in the RocA X-ray crystal structure (PDB: 5ZC9).^41^

Computational docking programs such as Glide perform quite well at estimating the enthalpic contributions to ligand binding, but entropic effects are significantly more difficult to capture due to the complex inherent molecular dynamics and need to consider entropy changes in the protein and bulk solvent environment, in addition to changes in ligand entropy.^42–46^ As such, entropic contributions are typically approximated in scoring functions at a low level of accuracy. For example, the standard precision Glide scoring function, which employs a modified version of the ChemScore scoring function, employs a rotatable bond term as an indirect approximation of the entropic penalties arising from conformational restriction of the ligand.^31,47^ More recent efforts to improve the precision of Glide scoring (“Glide XP” scoring) have attempted to address these concerns, but have not achieved significant advancements in this area.^47^ Importantly, since the fused cyclopenta[*b*]benzofuran ring system is entirely comprised of “non-rotatable” bonds, we assume that any reduction in confromational entropy caused by further rigidification of this system would likely not be adequately captured by Glide’s scoring functions. To better address this consideration, we instead sought to systematically examine and compare the conformational landscapes of **CMLD012824** and RocA to characterize how this apparent rigidification impacts the general eIF4A/RNA-binding pharmacophore of rocaglates.

We targeted a library of DFT-optimized conformers for each ligand using the established process of first deploying molecular mechanics to generate a library of unique low-energy conformers via fast and computationally inexpensive force field calculations, followed by more computationally-intensive density functional theory (DFT) calculations on these conformers to arrive at more accurate minimized geometries and computed energies.^48^ During the initial Macromodel conformational search stage, we noted a conspicuous absence of predicted RocA conformers in the eIF4A-RNA-bound cyclopenta[*b*]benzofuran conformation when using the best-parameterized force fields (OPLS4 and OPLS_2005).^49–51^ For both force fields, the closest all-atom RMSD from the bound conformation found among the conformational search output was >1.3 Å. This outcome was particularly concerning given prior computational studies on RocA, which identified a DFT-optimized lowest-energy conformer reported to have high conformational similarity to the X-ray liganded structure. Notably, force fields with lower quality parameters (MM3, MM2, AMBER, OPLS, MMFF and MMFFs) all produced at least one conformer with RMSD <0.3 Å compared to the X-ray bound conformation (**Supplementary Table S1**).^41,52–55^ To resolve this issue, the OPLS4 conformational search was repeated with a slightly extended energy window for retaining structures, increasing the 21.0 kJ/mol (5.02 kcal/mol) cutoff to 25.0 kJ/mol (5.97 kcal/mol), in line with recommendations by Sharapa and coworkers.^56^ This resulted in a library of 24 unique RocA conformers, four of which closely resembled the X-ray bound RocA ligand (heavy-atom RMSD ≤1 Å).

Examination of the cyclopenta[*b*]benzofuran ring system conformation in each of the RocA conformers revealed that the benzofuran region is predominantly planar/flat, and that core differences are driven by pseudorotation of the fused cyclopentane ring (**Figure 6A**). The conformers were sub-classified based on their cyclopentane envelope (or slightly twisted envelope) conformations, using a classification system wherein a different cyclopentane ring carbon assumes the “exo” position, pointed either above (“up”) or below (“down”) the plane of the benzofuran ring (according to the perspective depicted in **Figure 6A**). Three core conformational clusters were identified for RocA (**Figure 6B** and **Supplementary Figure S4**). The first cluster, which we named **RocA I**, was comprised of the 16 lowest energy conformers as predicted by force field calculations. In this core conformation, we observed the *C*1 carbon to adopt the *exo* position, positioned “up” relative to the benzofuran (*C*1-exo_up_). A second cluster of four conformers, which showed a *C*2-exo_down_ conformation, was assigned as **RocA II**. Lastly, the aforementioned subpopulation of four conformers binned as **RocA III** showed a cyclopentane envelope with the *C*3-exo_down_ conformation, in excellent alignment with the bound RocA ligand from the published X-ray structure (PDB: 5ZC9, **Figure 6C, Supplementary Figure S4** and **Supplementary Figure S5**), but were predicted by force-field calculations to be significantly higher in energy (+∼5.7 kJ/mol) than the lowest-energy **RocA I** conformer.

**Figure 6.**
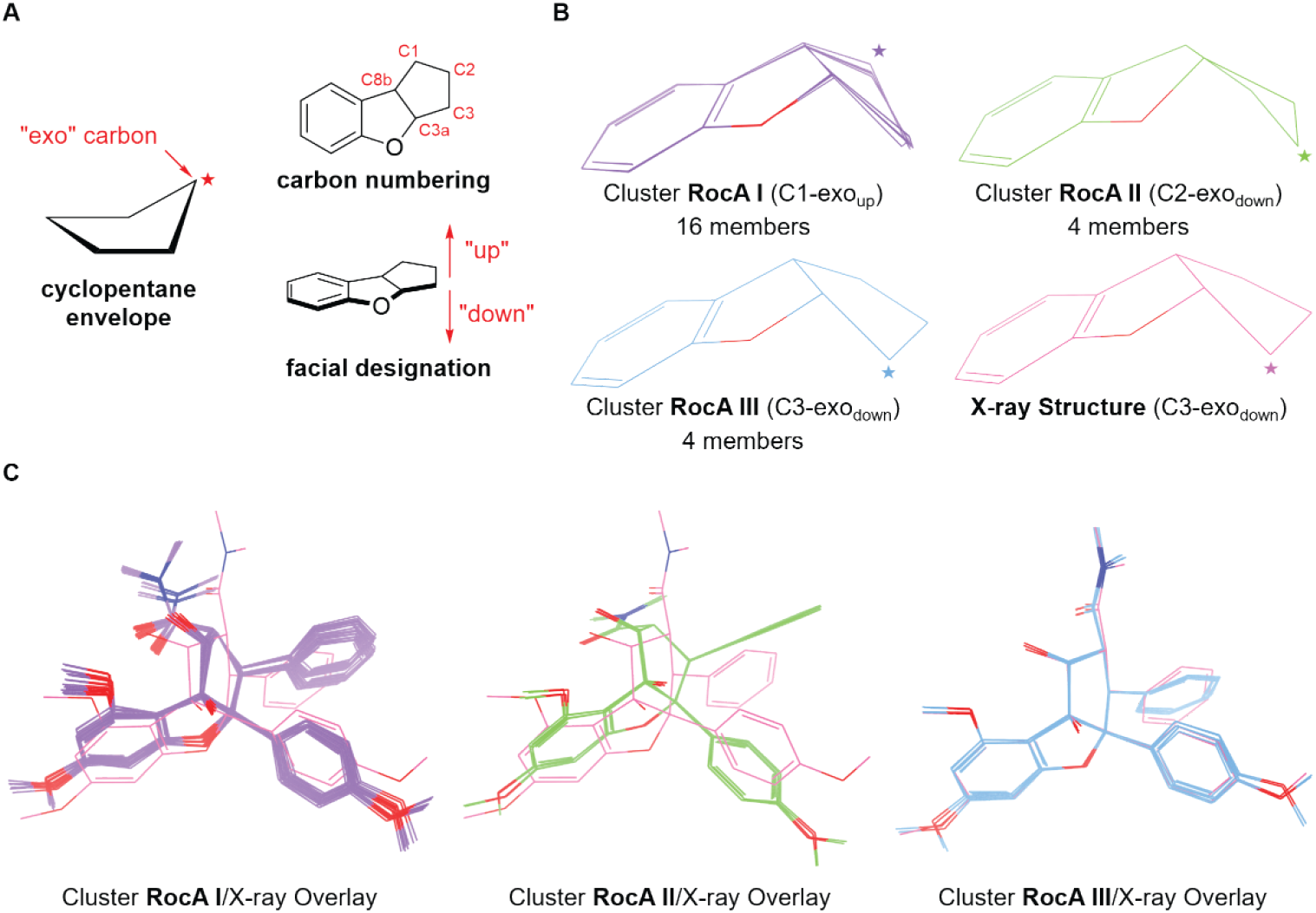
Conformational analysis of force-field computed RocA conformers. **(A)** Guide to nomenclature and numbering convention for cyclopenta[*b*]benzofuran core conformational classes based on the cyclopentane envelope conformation. **(B)** Overlays of each conformational cluster with only core atoms depicted, with the RocA X-ray bound core also depicted for comparison. For each depicted core conformation, the exo carbon is denoted with a star symbol. **(C)** All-atom overlays of each conformational cluster for RocA with eIF4A-bound RocA from the X-ray structure (PDB: 5ZC9; *pink*).

This Macromodel conformational search analysis was extended to the ADR **CMLD012824**, again saving all unique conformers within a 25 kJ/mol (5.98 kcal/mol) energy window. This search produced 181 unique conformers that were similarly analyzed and subdivided into six unique core conformational clusters (**Figure 7A** and **Supplementary Figures S6** and **S7**). Cluster **ADR I** contained 44 unique conformers, each of which had a *C*2-exo_down_ conformation (analogous to cluster **RocA II**). Cluster **ADR II**, initially populated by 43 unique conformers, showed the same *C*3-exo_down_ conformation observed in the complexed RocA/**CMLD012824** structures (Figure CB), and was conformationally analogous to **RocA III**. Cluster **ADR III** (*C*3-exo_up_) contained 76 members and represents a cyclopentane ring inversion from cluster **ADR II**. Cluster **ADR IV** (*C*2-exo_up_), which contained 15 members, showed the second-lowest average RMSD to eIF4A-complexed **CMLD012824**, but with a slight pseudorotation of the twisted envelope. Lastly, three additional high-energy conformers were assigned to clusters **ADR V** and **ADR VI**, with unique envelope conformations placing the *C*8b and *C*3a ring fusion carbons at the exo position, respectively (*C*8b-exo_up_ and *C*3a-exo_up_). Of note, no ADRs adopted the *C*1-exo_up_ conformation that predominated the RocA conformer library (cluster **RocA I**), likely due to the conformational strain imposed by the imidazole ring fusion at *C*1 blocking the required pseudorotation.

**Figure 7.**
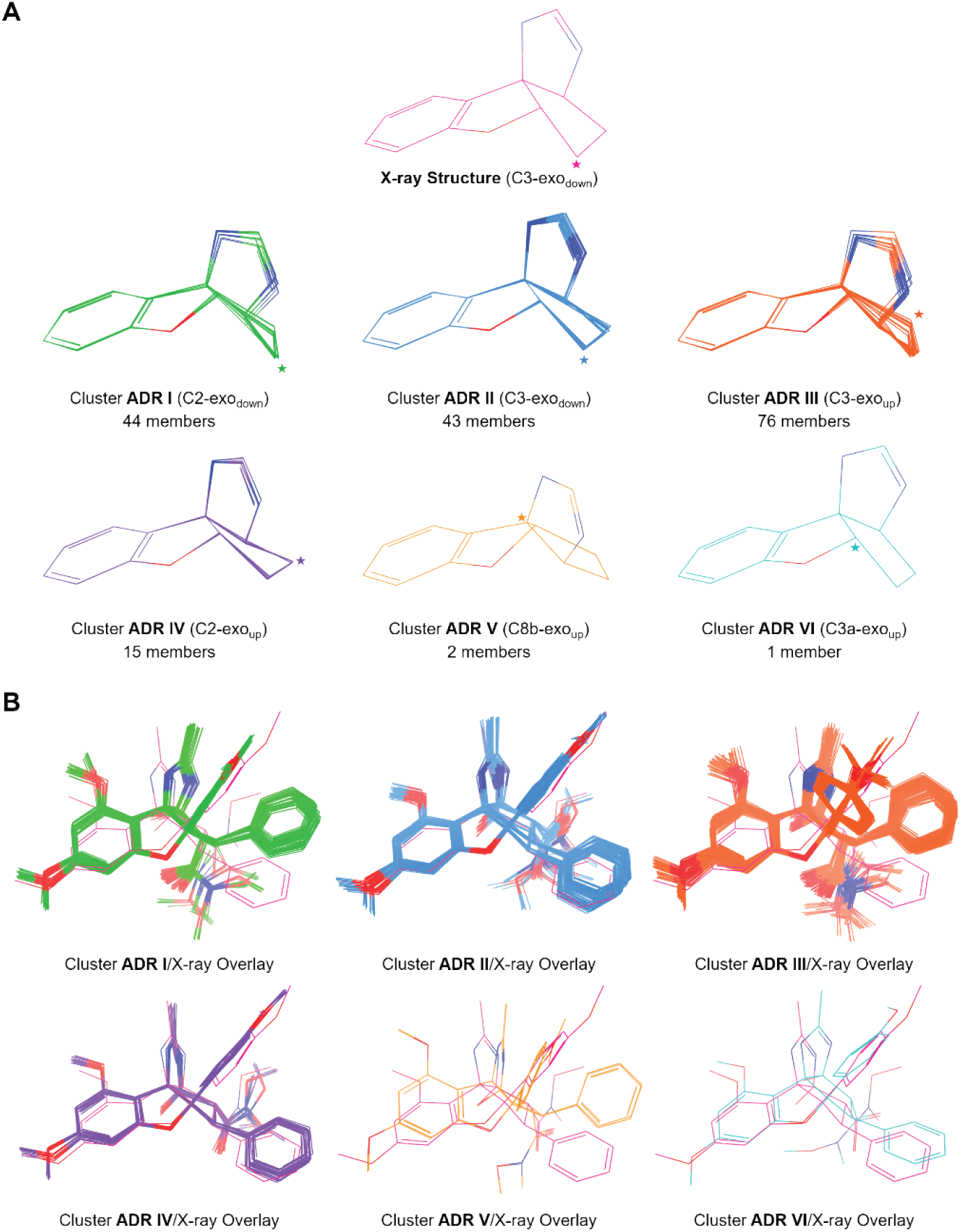
Conformational analysis of force-field computed CMLD012824 conformers. **(A)** Overlays of each conformational cluster with only core atoms depicted, with the **CMLD012824** core from the liganded X-ray structure depicted for comparison. For each depicted core conformation, the exo carbon is denoted with a star symbol. **(B)** All-atom overlays of each conformational cluster for **CMLD012824**, aligned with eIF4A-bound **CMLD012824** from the X-ray structure (*pink*).

We next applied density functional theory (DFT) calculations to further refine the geometry of each conformer and to understand how the energetics of each conformational subpopulation compare to one another at a higher level of computational accuracy. Each conformer was geometrically optimized in Jaguar using B3LYP-D3/6-31g**, followed by single-point energy calculations using M06-2X/6-31g++** to predict relative gas phase energies.^57,58^ For both ligands, the geometry optimizations slightly altered the distribution of conformers across their established subpopulations (**Supplementary Tables S2 and S3**), and also led to the identification of one new sparsely populated, high-energy core cluster for each ligand (**Supplementary Figures S8 - S11**). Specifically, in the case of RocA, a new high-energy subpopulation (**RocA IV, Supplementary Figures S8** and **S9**) with four members was identified with a C8b-exo_down_ conformation that, while distinct from cluster **RocA III**, more closely mimicked the bound conformation and projection of the B- and C-aromatic rings than clusters **RocA I** and **RocA II** (**Supplementary Figure S9**). For **CMLD012824**, three high-energy conformers (relative energies ≥ +11.7 kcal/mol) converged on a new geometry (**ADR VII**) with a C3a-exo_down_ conformation (**Supplementary Figures S10 and S11**).

To better visualize the energy landscape of the DFT-optimized conformer libraries, we plotted the heavy-atom RMSD from each conformer’s respective crystallographically-determined ligand conformation against its relative M06-2X/6-31g++** gas phase energy (**Figure 8**). The analysis primarily focused on the conformers within a relative gas-phase energy window of 0-2 kcal/mol (**Figure 8A and 8B**, dashed gray line) based on the expectation that conformations with relative energies >2 kcal/mol from the lowest-energy conformer are not expected to be substantially populated.^59–61^ Nonetheless, for **CMLD012824** (**Figure 8A**), we noted that a wider energy range (spanning from 0 to +3.5 kcal/mol) was exclusively populated with core conformation **ADR II**, the conformation with highest similarity to X-ray structure of bound **CMLD012824**. Further, all members of cluster **ADR II** showed a heavy atom RMSD <1.5 Å from that of the X-ray structure of bound ligand, with higher RMSDs (>1.0 Å, ascribable to various rotamers of the C2-tethered Weinreb amide) generally correlating with higher computed energies. Together, these observations strongly support the premise that **CMLD012824** predominantly exists in a conformational state that is inherently pre-organized to clamp eIF4A onto RNA.

**Figure 8.**
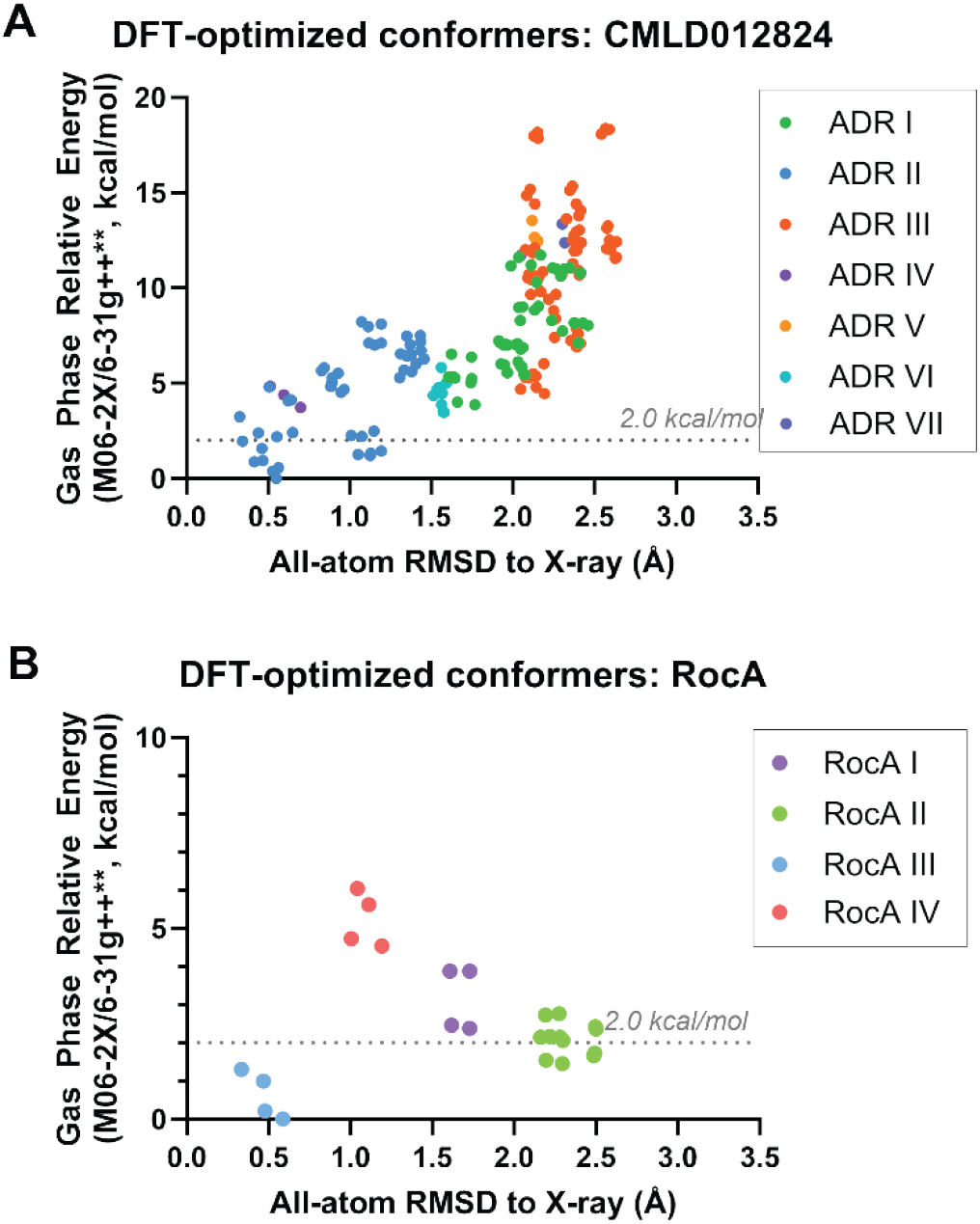
Energetic landscape of CMLD012824/RocA conformer classes. **(A)** Scatter plot of the heavy atom RMSD (in Å) from the **CMLD012824** X-ray structure vs the DFT-computed gas phase relative energy (M06-2X/6-31g++**, in kcal/mol) for each unique geometry-optimized **CMLD012824** conformer. Data points are colored according to conformational cluster. Gray dashed line indicates the upper boundary of the 2.0 kcal/mol energy window for substantially populated conformational states at room temperature. **(B)** Scatter plot of the heavy-atom RMSD (in Å) from the RocA X-ray structure vs the DFT-computed gas phase relative energy (M06-2X/6-31g++**, in kcal/mol) for each unique geometry-optimized RocA conformer. Data points are colored according to conformational cluster and gray dashed line defined as in **(A)**.

Extending similar analysis to RocA (**Figure 8B**) we found that in contrast to **CMLD012824**, two distinct clusters (**RocA II** and **RocA III**) sat within the 0-2 kcal/mol energy window. Based on force-field derived energies, we had considered the possibility that class **RocA III**, which most resembles the eIF4A-RNA-bound ligand (all RMSDs ∼0.5 Å), represented a highly strained conformation. However, this turned out not to be the case, as this cluster was determined by DFT to be the lowest energy conformer class. This outcome is both consistent with prior findings in Zotatifin development, and is perhaps unsurprising due to the known poor correlation between force field and DFT-derived energies.^41,60^ The second lowest-energy cluster, **RocA II**, shows a significant conformational deviation from the X-ray bound conformation (all RMSDs >2 Å). Although cluster **RocA II** is slightly higher in computed energy than cluster **RocA III**, we cannot exclude the possibility that clusters **RocA III** and **RocA II** are degenerate, given to the inherent ∼2 kcal/mol accuracy limits of DFT.^61,62^ Importantly, cluster **RocA II**, which bears a C2-exo_down_ conformation that was not observed for any ADR, represents a reasonably populated low-energy core conformational state that would be disallowed upon eIF4A-RNA binding. It is also worth noting that a third cluster (**RocA I**) was also found in the expanded energy window of 2 - 3.5 kcal/mol. While the computed relative energies suggest that this conformation may not be populated to a significant extent at room temperature, the presence of a third conformational cluster within the same energy window that was exclusively populated with pre-organized (**ADR II**) conformation of **CMLD012824** further underscores the conformational plasticity of the RocA core relative to the more rigidified ADR.

Collectively, these calculations strongly support our hypothesis that the fused amidine ring imparts significant conformational rigidification of the rocaglate scaffold, locking the core in a productive conformation for eIF4A-RNA binding. This preorganization ensures that **CMLD012824** likely engages eIF4A and RNA without experiencing a significant loss in entropy upon binding, contrasting more conformationally flexible, non-amidino rocaglate ligands such as RocA. This phenomenon likely contributes to the observed enhanced RNA clamping ability and translation inhibition potency of ADR **CMLD012824** relative to non-amidino rocaglates.

## Conclusions

Rocaglates have emerged as a unique class of compounds of considerable interest in drug discovery, possessing a unique quality as small molecules with sequence-selective translational repression. Additional work is needed to expand our understanding of these compounds to develop them into potent anticancer and anti-infectious agents. The present investigation represents an important step in that direction, with the determination of the X-ray crystal structure of the amidino-rocaglate (ADR) **CMLD012824**, bound to the DEAD box helicase eIF4A1, a compound which had been shown to exhibit very strong “clamping” of eIF4A1 to a polypurine RNA probe. In this study, we identified the conserved interactions of rocaglates and ADRs with a polypurine RNA sequence and eIF4A1. Our computational docking experiments align with these structural observations that suggest that both RocA and **CMLD012824** engage RNA in a similar way and have similar enthalpic gains upon binding, prompting an investigation into the entropic determinants of binding for each of these rocaglate subtypes. DFT-optimized computational modeling illustrated a difference in the distribution of conformers of RocA and **CMLD012824**. Gas-phase energy calculations of these different conformer subpopulations were categorized into subpopulations for both RocA and **CMLD012824**, with the ADR having a single subpopulation within a ∼2 kcal/mol window of the experimentally-determined conformer from the X-ray structure, while RocA existed in two unique conformers within this window. These findings suggest that the ADR, which possessed a unique amidine ring fusion on the rocaglate scaffold, was inherently preorganized in a conformation capable of clamping eIF4A1 onto polypurine RNA segments. Our study clarifies how the rigidifying effect of the ADR scaffold improves “clamping” and thereby potency which will be helpful for future development of ADRs and congeners as potential anti-neoplastic, antiviral, and antiparasitic agents.

## Experimental Section

### Construct Design and Protein Expression

A sequence encoding residues 21-406 of eIF4A1 with codons optimized for expression in *E. coli* (Twist BioSciences) was cloned using Gibson Assembly into a pTB146 vector to include an N-terminal 6x histidine tag followed by a Small Ubiquitin-like Modifier (SUMO) tag. The truncation of the first 20 amino acids of eIF4A1 was based on the lack of electron density for these residues in the previously published structure of eIF4A1:RocA (5ZC9). The resulting pTB146-His_6_-SUMO-eIF4A1(21-406) was transformed into BL21(DE3) cells (New England Biolabs), plated, and incubated overnight. Single colonies were isolated and used to inoculate small “starter” cultures of Luria Broth containing 100 μg/mL carbenicillin and grown overnight for ca. 12-15 h at 37 °C. These cultures were used to inoculate large-scale growths, grown at 37 °C until OD_600_ = 0.6-0.8. Protein expression was induced with IPTG added to a final concentration of 1 mM. Flasks were incubated overnight for ca. 18 h at 16 °C, whereupon cells were harvested by centrifugation at 4000*g* for 10 min. The cell pellet was stored at -80°C.

### Protein Purification

eIF4A1(21-406) was purified as previously described with the following modifications.^63^ Clarified lysate was applied to HisPur™ Ni-NTA Resin in a gravity column instead of using an FPLC. Protein was eluted in a single step using 20 mL of a buffer containing 262.5 mM imidazole. Eluted protein was dialyzed overnight together with SUMO Protease into a buffer lacking imidazole to cleave purification tags followed by subtractive IMAC (SUMO Protease prepped in house according to a published protocol).^64^ Size-exclusion chromatography (SEC) on a 26 mm × 600 mm HiPrep 26/60 Sephacryl S-300 HR column (Cytiva), equilibrated in 50 mM EPPS-KOH pH 8.0, 100 mM KCl, 5 mM MgCL_2_, 1 mM DTT, and 0.1 mM EDTA was used to achieve a homogeneity (evaluated by SDS-PAGE and >80% monodispersity in dynamic light scattering) sufficient for crystallization experiments. Amicon Ultra centrifugal filter devices (30 kDa cutoff) were used to concentrate the protein before and after SEC. SEC-purified fractions were concentrated to between 8 and 15 mg/mL, whereupon glycerol was added to a final glycerol concentration of 10%, and aliquots flash-frozen in liquid nitrogen and stored at -80 °C.

### Chemical Synthesis

The chiral, enantioenriched ADR **CMLD012824** was synthesized by condensation of a rocaglate enol tosylate precursor (>99% e.e.) with methyl amidine according to our literature procedure.^23^ The compound employed in this study possessed >95% purity as determined by UPLC-MS-ELSD analysis.

### Crystallization

Purified eIF4A1(21-406) (5 - 15 mg/mL) was mixed with super-stoichiometric (1.5x) amounts of **CMLD012824**, nonhydrolyzable ATP-analog AMPPNP (5x), and poly (AG)_5_ RNA (3x). AMPPNP (Millapore Sigma) and poly (AG)_5_ RNA (Horizon Discovery/Dharmacon) were purchased commercially. Protein cocktail was mixed with well solution in various proportions to yield a protein concentration of 10 mg/mL after equilibration. Vapor diffusion with sitting-drop geometry at 17 °C was utilized. Initial cubic crystals, smaller than 25 microns, were identified from a sparse matrix screen (Hampton Research PEG/Ion) in a condition comprised of 0.2 M ammonium acetate and 20% w/v PEG 3350. Crystal optimization strategies included improving protein purification quality through more stringent selection of purified fractions and varying the precipitant polymer. The final condition comprised 0.2 M ammonium acetate and 8% PEG20000 and 10% Hampton Silver Bullet Reagent B5, a crystallization additive cocktail comprised of 0.33% w/v 2,7-naphthalenedisulfonic acid disodium salt, 0.33% w/v azelaic acid, 0.33% w/v trans-cinnamic acid, 0.02 M HEPES sodium pH 6.8. *N*.*B*. The crystal used to collect the final dataset was grown from a condition including the Hampton Silver Bullet Reagent B5, however several crystals were harvested from wells containing other Silver Bullet additives and diffracted to similar resolution in the same space group. It does not appear that the Silver Bullets altered crystallization or diffraction characteristics in this experiment.

### Data Collection and Structure Refinement

Crystal harvesting/cryo-protection solutions were prepared by reproducing the well condition with the addition of 10% glycerol. Crystals were harvested from the mother liquor and incubated in the cryoprotection solution for 30 s - 3 minutes prior to flash-cooling in liquid nitrogen at 100 K. X-ray diffraction data was collected at the Brookhaven National Laboratory at the National Synchrotron Light Source II (Upton, NY) on beamline 17-ID-1 (AMX) outfitted with an Eiger 9M detector. Diffraction data were collected under nitrogen gas cooled to 100 K and were processed, integrated, and scaled using XDS.^65^ Initial phases were calculated using the previously determined structure of eIF4A1 in complex with RocA, ploy (AG)_5_, and AMPPNP (PDB: 5ZC9) with small molecule ligands and waters removed as a search model for molecular replacement in PHENIX Phaser-MR.^66^ Multiple rounds of refinement in PHENIX Refine using individual B-factors, TLS parameters, and addition of waters, along with manual model building were used in iterative rounds.^67^ Ligands were positioned in electron density manually using Coot and validated using Fo − Fc omit and Polder electron density maps in PHENIX.^68^ All images were generated in PyMOL.^69^

### Computational Docking

RocA and **CMLD012824** ligands were prepared for docking using the LigPrep application in the Maestro software environment (Version 2024-2, Schrödinger LLC). Likewise, eIF4A/RNA receptor structures were prepared from the 5ZC9 X-ray structure, and the ADR-bound X-ray structure reported herein using the default Protein Preparation Workflow in Maestro. The protein preparation protocol involved structure pre-processing, hydrogen-bond optimization, and restrained minimization (S-OPLS force field, hydrogen atoms freely minimized and heavy atoms minimized to RMSD 0.3 Å, and removal of water >5Å from heavy atoms). Docking grids were then generated from the prepared structures, sized and centered at the existing rocaglate/ADR ligand. Glide docking of each ligand into its respective receptor grid was then performed using default settings at standard precision (SP), with five output poses produced per ligand. The top scored pose (Glide G_scores_) for each compound was selected for analysis.

### Macromodel Conformational Searches

Conformer libraries for RocA and **CMLD012824** were generated using the MacroModel Conformational Search tool (force field: OPLS4, method: torsional sampling/MCMM, solvent: water, torsional sampling: enhanced, energy window for saving structure: 25 kJ/mol in the Maestro software environment (Version 2024-2, Schrödinger LLC). Conformer libraries were manually inspected to remove redundant conformers and binned according to core conformation with the aid of comparative overlays and RMSD calculation (either all-atom or select atom) using the “Superposition” tool in Maestro. Relative potential energies were calculated by subtracting the potential energy of each library member from the potential energy of the lowest-energy conformer in that library (for which relative potential energy = 0 kJ/mol).

### Jaguar DFT Calculations

The output conformer libraries from the Macromodel conformational search were subjected to DFT geometry optimizations in the gas phase using the Jaguar application in the Maestro software environment (Version 2024-2, Schrödinger LLC). Optimizations were performed at the B3LYP-D3 level of theory with the 6-31g** basis set. Each geometry-optimized structure was then subject to single point energy conformations using Jaguar at the MO6-2X level of theory with the 6-31g++** basis set. The total number of canonical orbitals employed were 831 for RocA, and 910 for **CMLD012824**. A heavy atom RMSD from the respective X-ray bound ligand for each conformer was calculated using the Superposition tool in the Maestro software environment. Scatter plots of heavy atom RMSD vs relative potential energy for all conformers were generated in Prism (Version 10.2.3, GraphPad, Inc.).

## Supporting information

Supplement containing Figs S1-S11, Tables S1-S3, Computational data and Supplemental References

## Associated Content

### Author Information

**James F. Conley** – *Department of Pharmacology, Boston University Medical School, 700 Albany St, Boston, MA 02215, United States*;

**Lauren E. Brown** – *Department of Chemistry and Center for Molecular Discovery (BU-CMD), Boston University, 590 Commonwealth Avenue, Boston, Massachusetts 02215, United States*;

**James H. McNeely** – *Department of Chemistry, Boston University, 590 Commonwealth Avenue, Boston, Massachusetts 02215, United States*;

**Jerry Pelletier** – *Department of Biochemistry, McGill University, 3655 Promenade Sir William Osler, Montreal, Quebec H3G 1Y6, Canada*;

**John A. Porco** – *Department of Chemistry and Center for Molecular Discovery (BU-CMD), Boston University, 590 Commonwealth Avenue, Boston, Massachusetts 02215, United States*;

**Karen N. Allen** – *Department of Chemistry, Boston University, 590 Commonwealth Avenue, Boston, Massachusetts 02215, United States*;

## Acknowledgements

.We thank the National Institutes of Health (NIH) (R35 GM118173 and U01 TR002625) for financial support. This research used the facilities of the AMX beamline of the National Synchrotron Light Source II, a U.S. Department of Energy (DOE) Office of Science User Facility operated for the DOE Office of Science by Brookhaven National Laboratory under Contract No. DE-SC0012704. The Center for BioMolecular Structure (CBMS) is primarily supported by the National Institutes of Health, National Institute of General Medical Sciences (NIGMS) through a Center Core P30 Grant (P30GM133893), and by the DOE Office of Biological and Environmental Research (KP1605010).

## Abbreviations Used

eIF4A1: eukaryotic Initiation Factor 4A-1;
RocA: rocaglamide A;
ADR: amidino-rocaglate;
DFT: density functional theory;
m7G: 5’-7-methylguanosine;
eIF: eukaryotic initiation factor;
UTR: 5’-untranslated region;
TEVp: tobacco etch virus protease;
SUMO: small ubiquitin-like modifier;
SEC: size exclusion chromatography;
LEC: lowest-energy conformer

