## Supplement containing Figs S1-S11, Tables S1-S3, Computational data and Supplemental References for "Structural Basis for the Improved RNA-Clamping of Amidino-Rocaglates to eIF4A1"

**Table of Contents**

**Supplementary Figure S1.** Rocaglate "family tree" .......... S2

**Supplementary Figure S2.** Structure of the eIF4A1-RNA-CMLD012824-AMPPNP asymmetric unit .......... S3

**Supplementary Figure S3.** Protomers of the eIF4A1-RNA-CMLD012824-AMPPNP asymmetric unit .......... S4

**Supplementary Figure S4.** Force field core conformations for RocA .......... S5

**Supplementary Figure S5.** Force field conformations for RocA .......... S6

**Supplementary Figure S6.** Force field core conformations for **CMLD012824** .......... S7

**Supplementary Figure S7.** Force field conformations for **CMLD012824** .......... S8

**Supplementary Figure S8.** DFT-optimized core conformations for RocA .......... S9

**Supplementary Figure S9.** DFT-optimized conformations for RocA .......... S10

**Supplementary Figure S10.** DFT-optimized core conformations for **CMLD012824** .......... S11

**Supplementary Figure S11.** DFT-optimized conformations for **CMLD012824** .......... S12

**Supplementary Table S1.** RocA conformational search results .......... S13

**Supplementary Table S2.** Summary of RocA conformational clusters .......... S13

**Supplementary Table S3.** Summary of **CMLD012824** conformational clusters .......... S14

**Computational Data** .......... S15

**Supplementary References** .......... S32

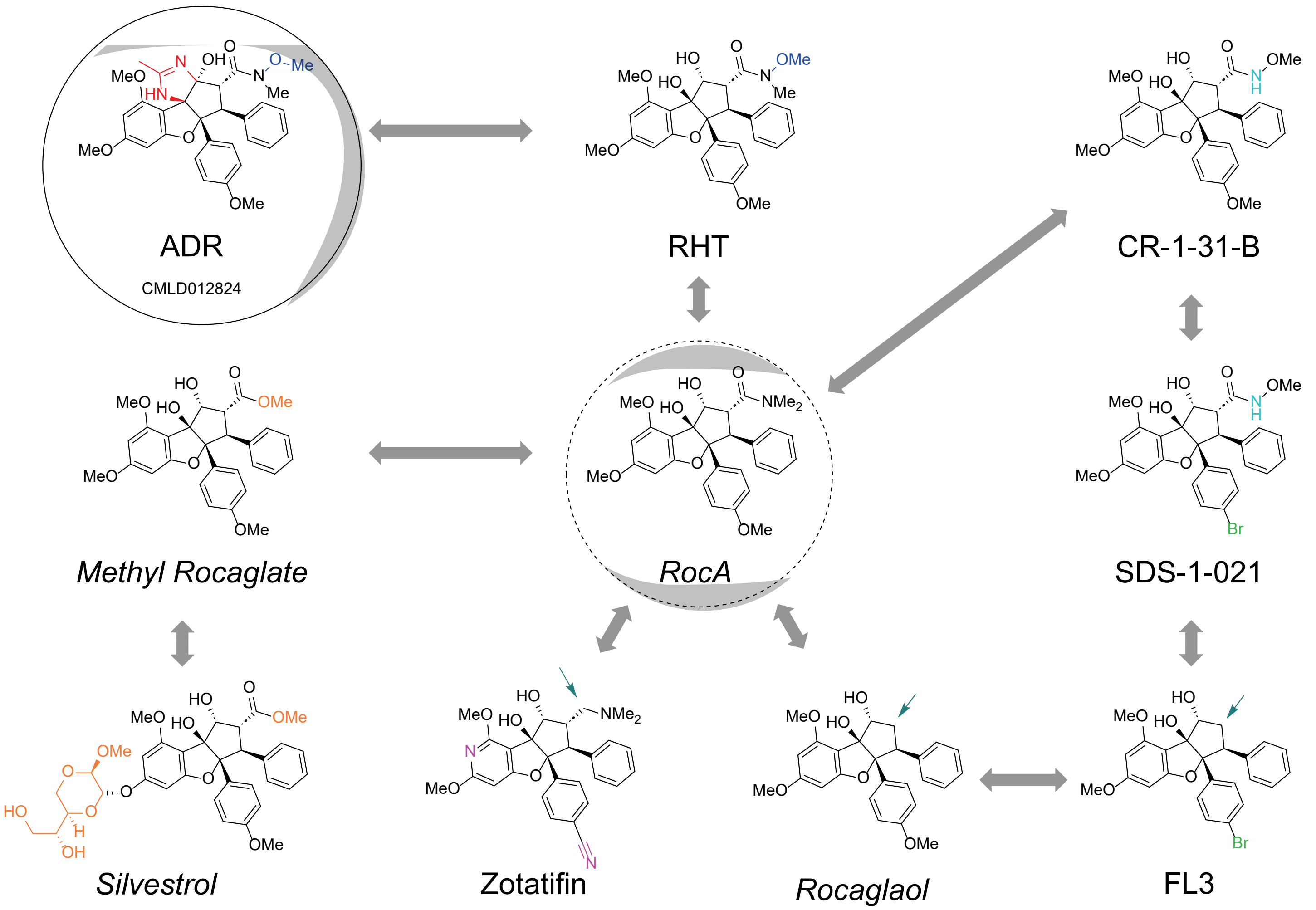


**Supplementary Figure S1.** Rocaglate "family tree." **Rocaglamide A** was the first isolated and characterized rocaglate, though many other rocaglates have since been isolated or synthesized. Gray lines added to emphasize similar compounds with minor differences. Small teal arrows denote when the differences result from the removal of functional groups. Compounds with italicized names were first isolated as natural products, while those with underlined names are of synthetic origin. **RocA** and **CMLD012824**, the two rocaglates highlighted in this manuscript are emphasized with dashed and filled circles, respectively.

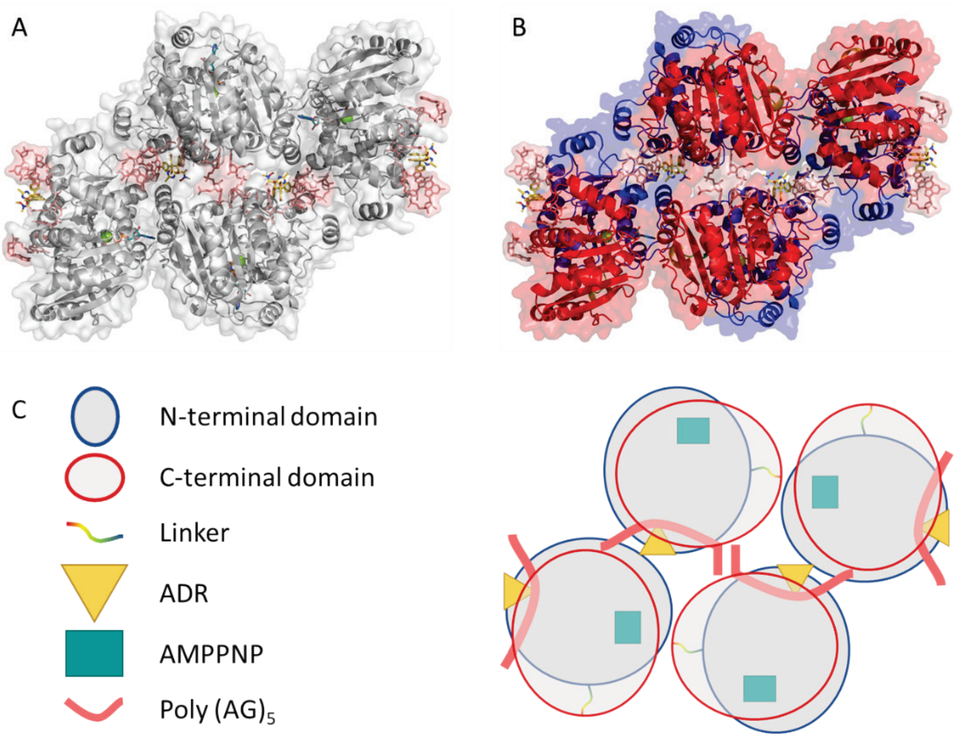


**Supplementary Figure S2. Intramolecular interactions of the eIF4A1-RNA-CMLD012824-AMPPNP asymmetric unit. (A)** eIF4A1 shown as off-white ribbon. Poly $(AG)_5$, CMLD012824, and AMPPNP shown as salmon, yellow, and teal sticks, respectively. **(B)** Panel A reproduced with eIF4A1 colored by domain: N-term, blue; C-term, red; linker (i.e. eIF4A1(230-243)), spectrum from blue to red. **(C)** Graphical representation of ASU (right) with legend (left) illustrating same side arrangement of N- and C- terminal domains of each protomer in the ASU, and unique conformation of central RNA oligonucleotides.

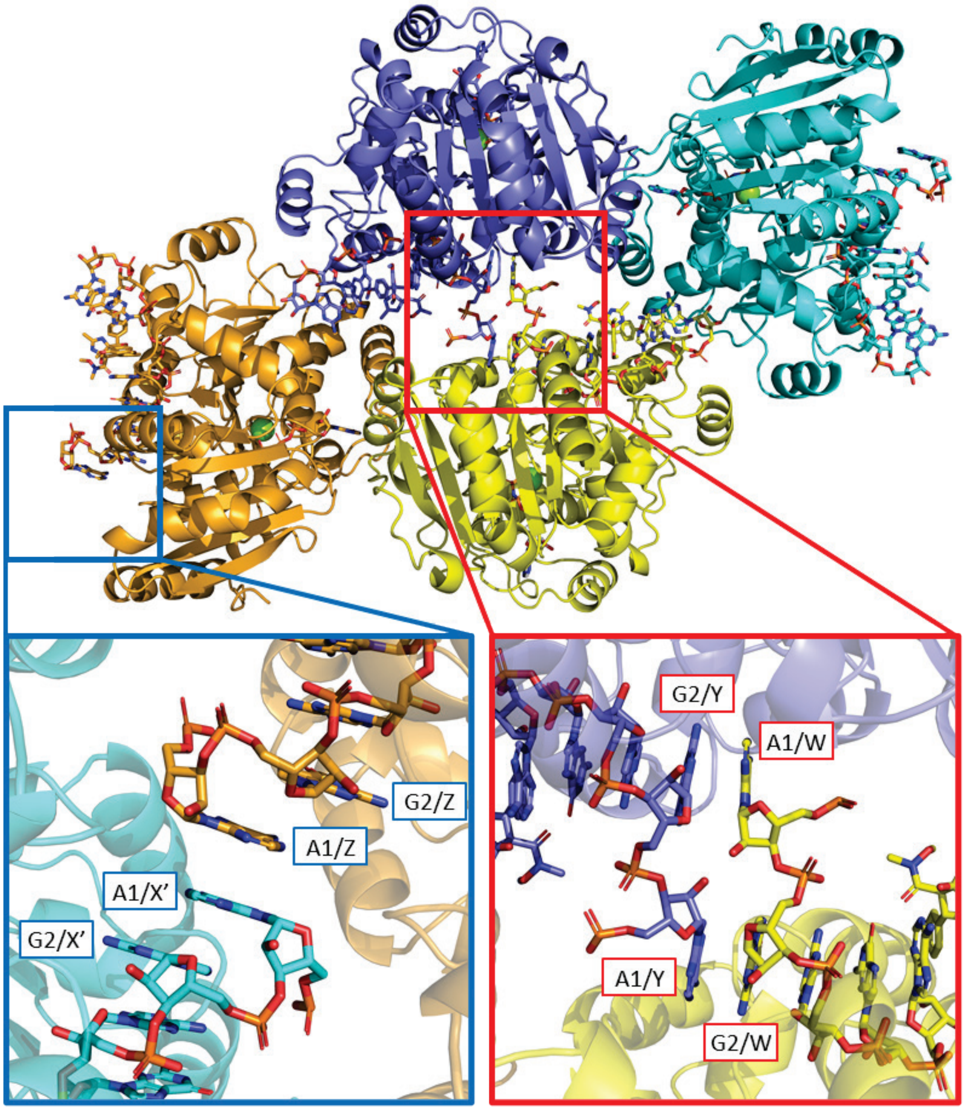


**Supplementary Figure S3. Differences in the protomers of the eIF4A1-RNA-CMLD012824-AMPPNP asymmetric unit.** Each quaternary complex is shown as a single color. Protein chains are termed A, B, C, D anticlockwise beginning with the bottom central quaternary complex, with corresponding associated RNA chains, W, X, Y, Z. π-stacking between nucleotides in chains X and Z is shown in expanded view in the blue box. π-stacking between nucleotides in chains W and Y is shown in expanded view in the red box, with its unique outward facing conformation of A1 from each chain.

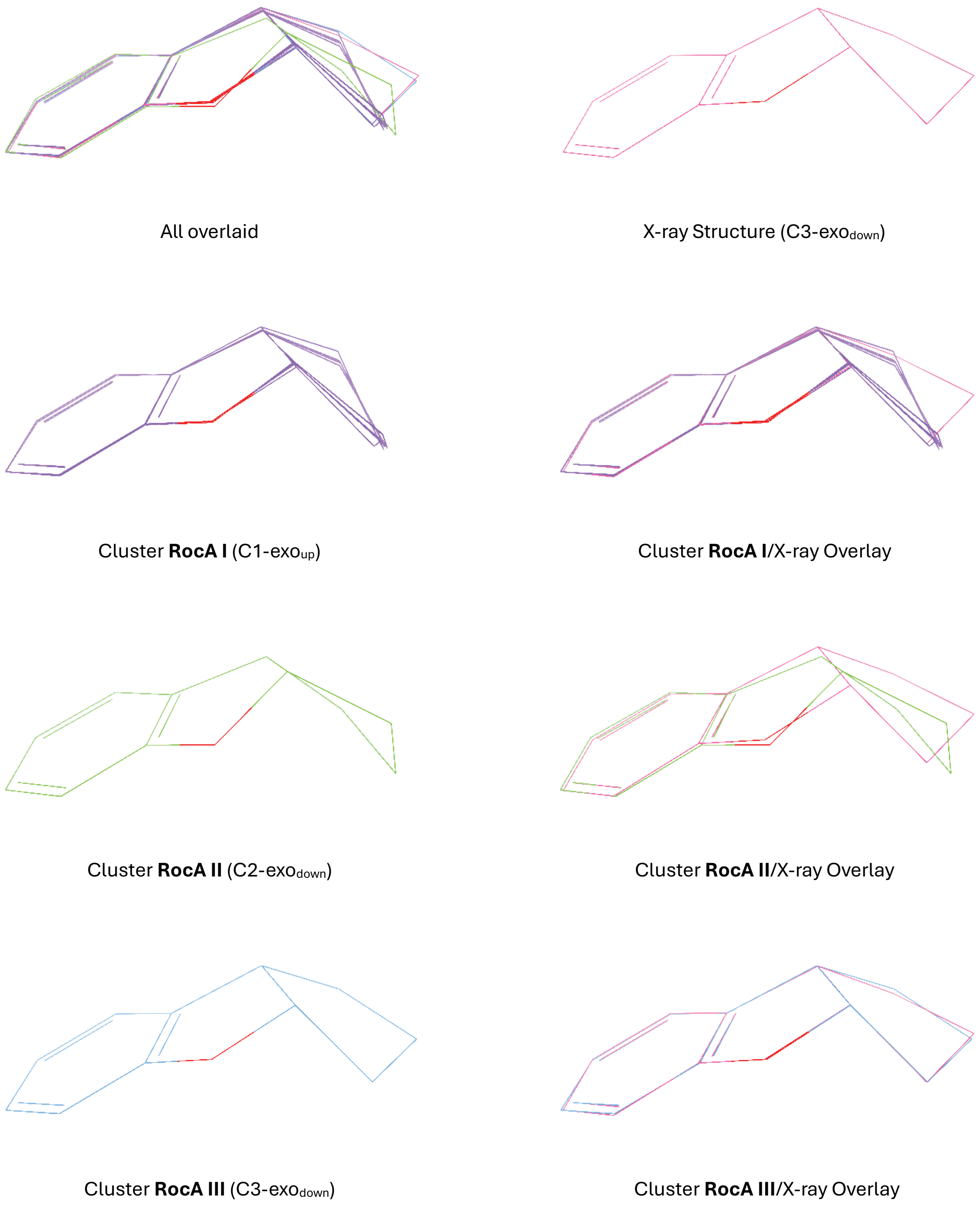


**Supplementary Figure S4.** Force field core conformations for RocA. Core conformational clusters for **RocA** found via Macromodel conformational search. Overlays were generated *via* core (cyclopenta[*b*]benzofuran) atom superposition (Maestro version 2024-2, Schrödinger LLC) of each output core conformer to the RocA X-ray core conformer.

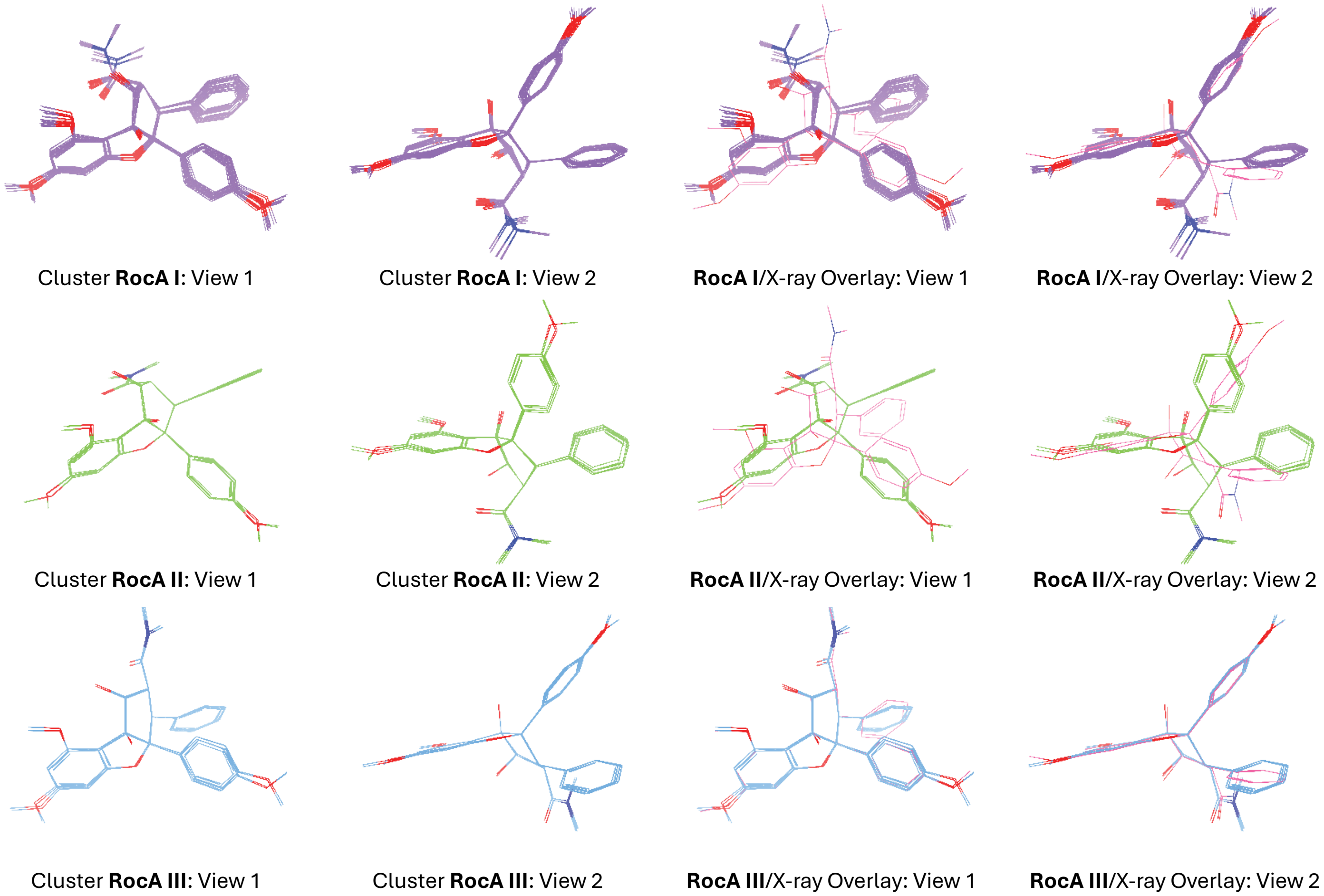


**Supplementary Figure S5.** Force field conformations for RocA. Conformational clusters for **RocA** found *via* Macromodel conformational search. Overlays were generated *via* heavy-atom superposition (Maestro version 2024-2, Schrödinger LLC) of each output conformer to the RocA X-ray core conformer (pink).

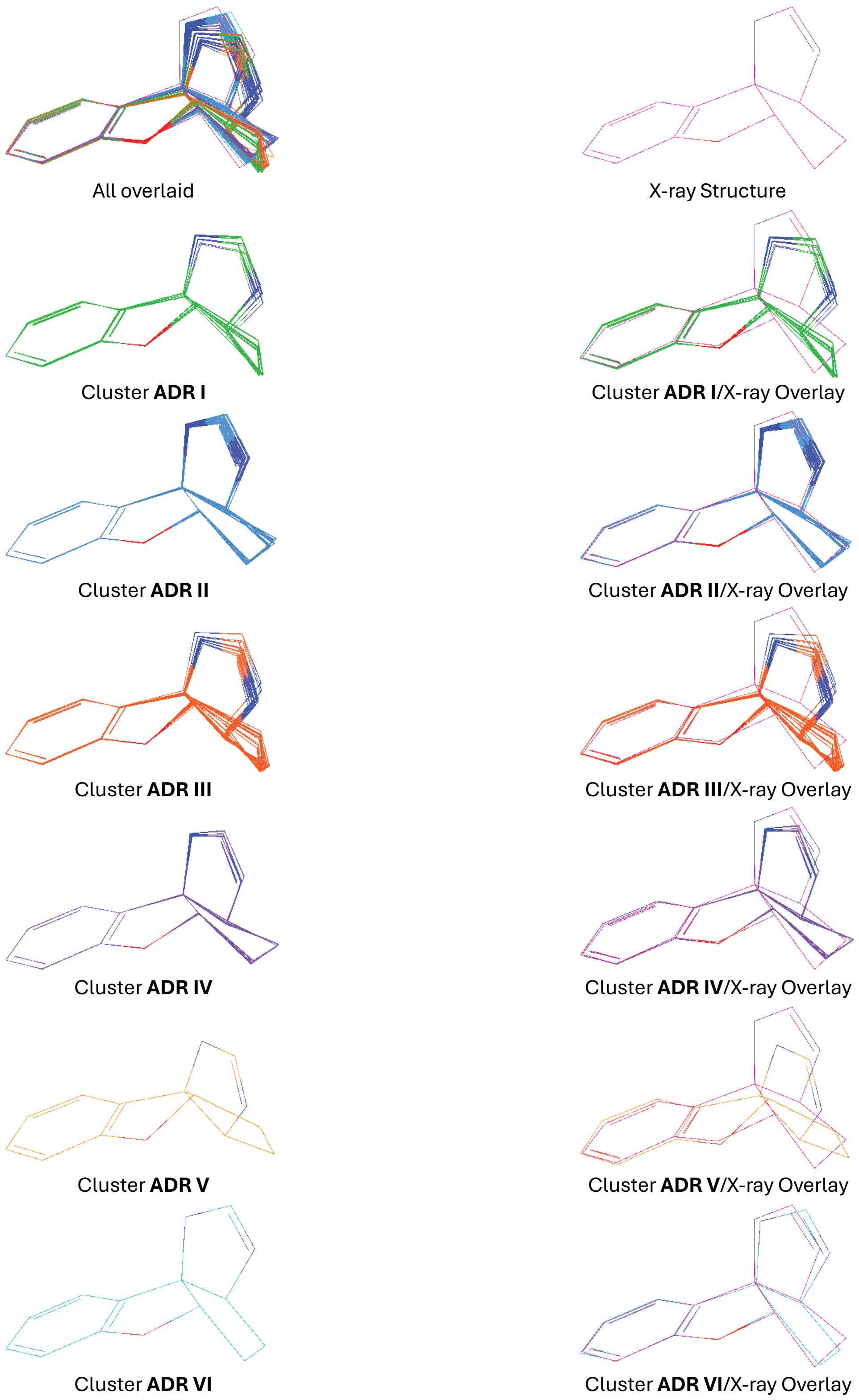


**Supplementary Figure S6.** Force field core conformations for **CMLD012824**. Core conformational clusters for **CMLD012824** found *via* Macromodel conformational search. Overlays were generated *via* core atom (tetrahydrobenzofurocyclopenta[1,2-*d*]imidazole) superposition (Maestro version 2024-2, Schrödinger LLC) of each output core conformer to the **CMLD012824** X-ray core conformer.

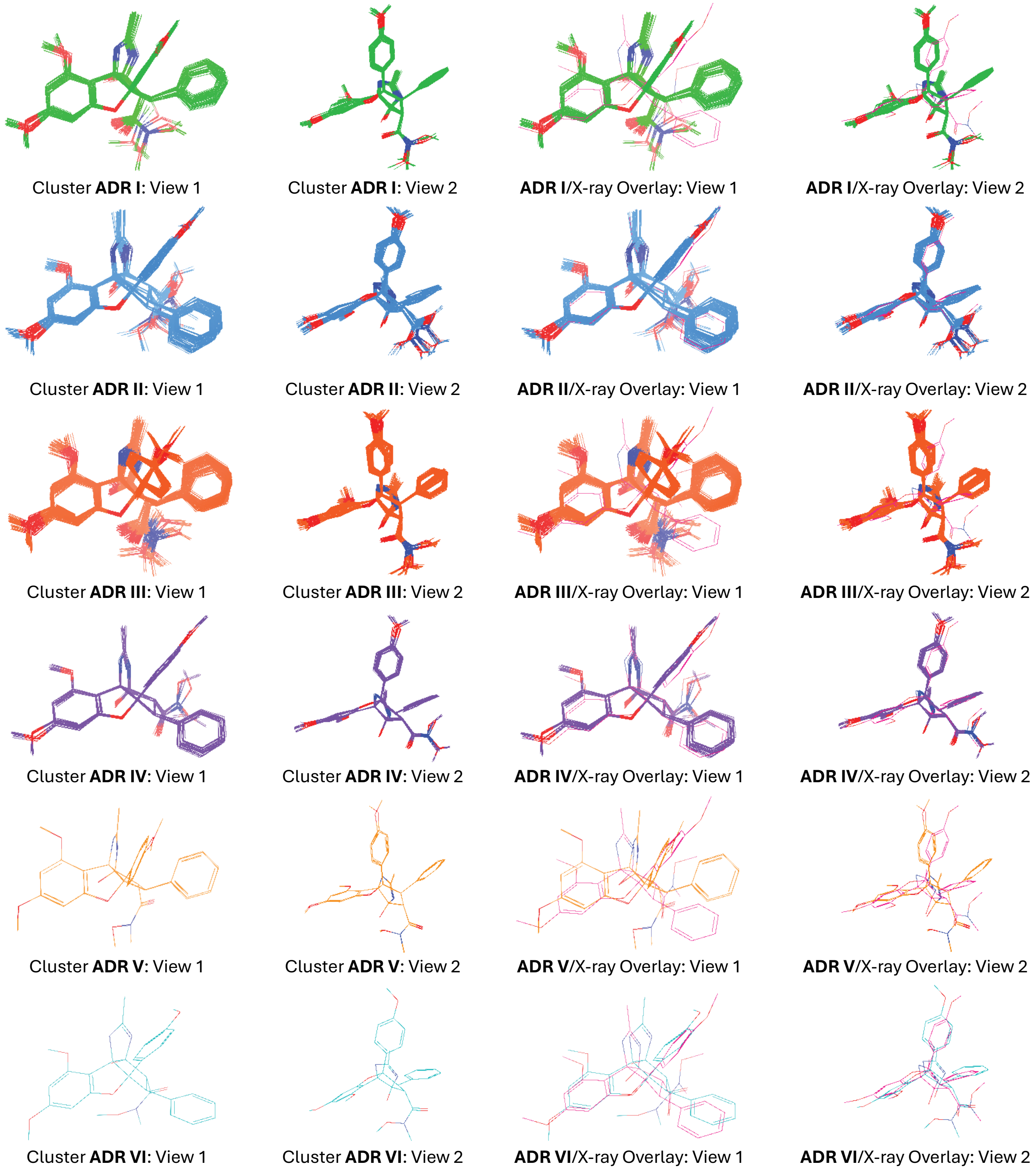


**Supplementary Figure S7.** Force field conformations for **CMLD012824**. Conformational clusters for **CMLD012824** found via Macromodel conformational search. Overlays were generated *via* heavy atom superposition (Maestro version 2024-2, Schrödinger LLC) of each output conformer to the **CMLD012824** X-ray conformer (pink).

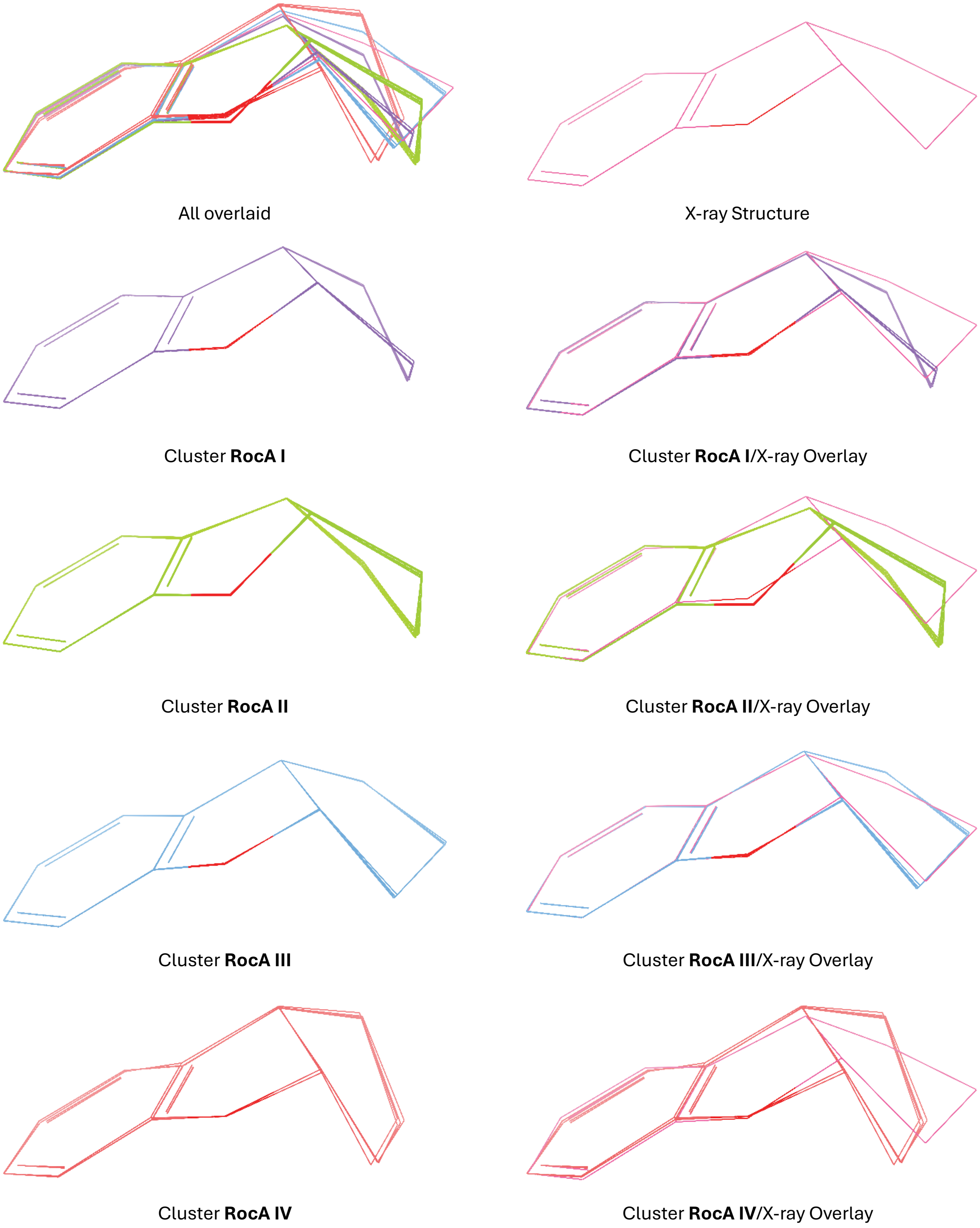


**Supplementary Figure S8.** DFT-optimized core conformations for RocA. Core conformational clusters for **RocA** after DFT (B3LYP-D3/6-31g**) geometry optimization. Overlays were generated *via* core (cyclopenta[*b*]benzofuran) atom superposition (Maestro version 2024-2, Schrödinger LLC) of each output conformer core to the RocA X-ray conformer core.

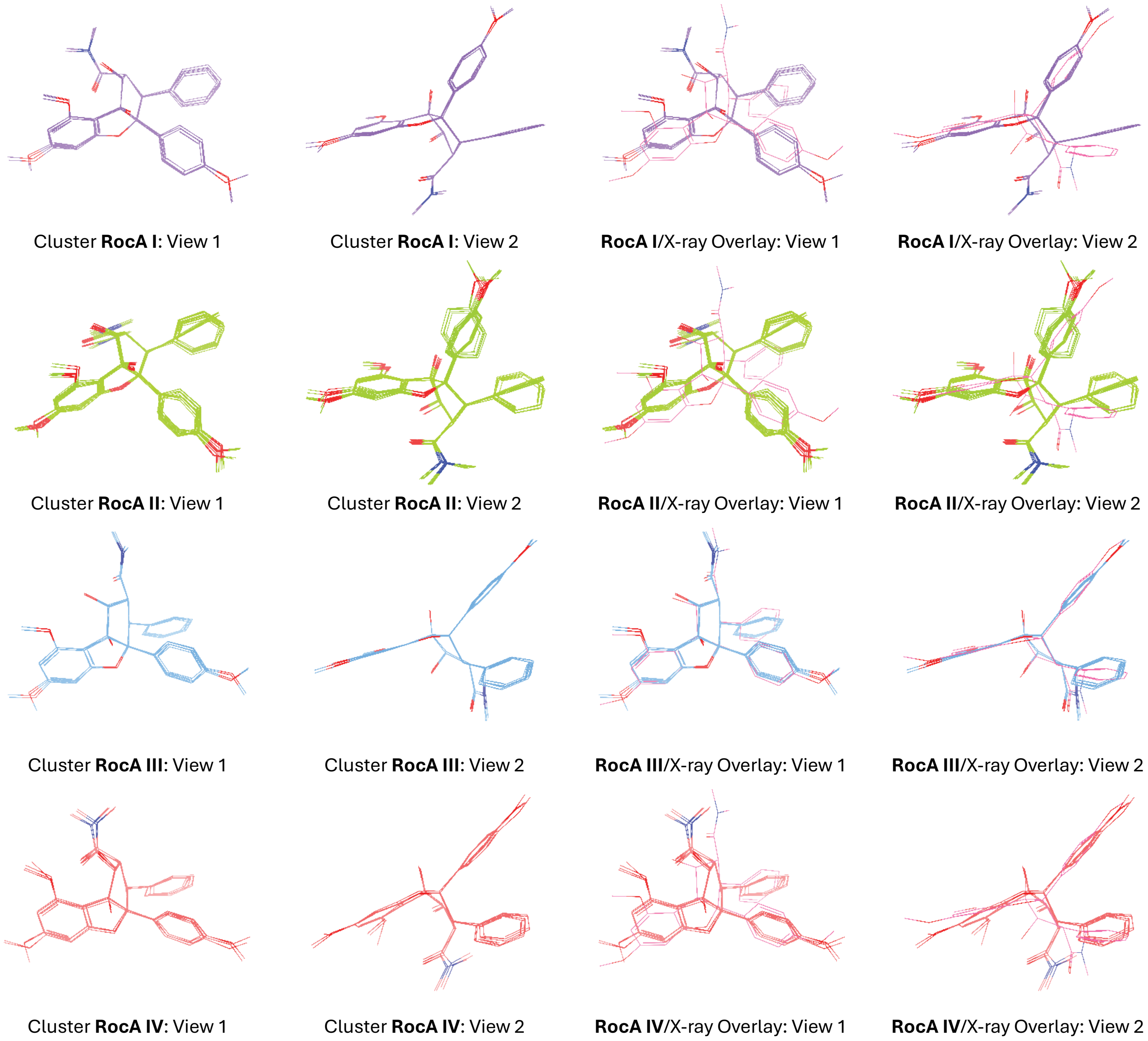


**Supplementary Figure S9.** DFT-optimized conformations for RocA. Conformational clusters for RocA after DFT (B3LYP-D3/6-31g**) geometry optimization. Overlays were generated *via* heavy atom superposition (Maestro version 2024-2, Schrödinger LLC) of each output conformer to the RocA X-ray core conformer (PDB: 5ZC9, pink).

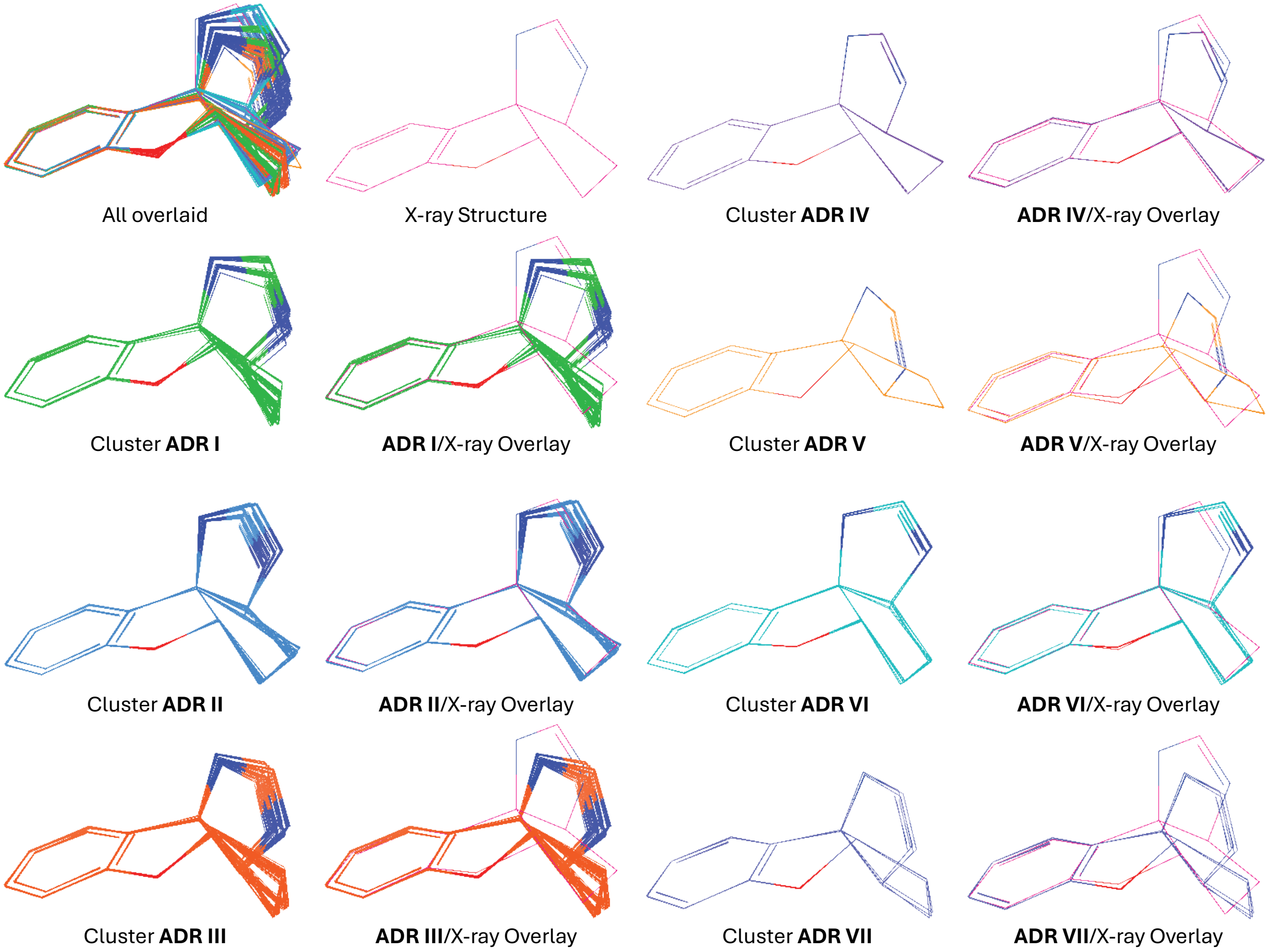


**Supplementary Figure S10.** DFT-optimized core conformations for **CMLD012824**. Core conformational clusters for **CMLD012824** after DFT (B3LYP-D3/6-31g**) geometry optimization. Overlays were generated *via* heavy atom superposition (Maestro version 2024-2, Schrödinger LLC) of each output conformer core to the **CMLD012824** X-ray conformer core.

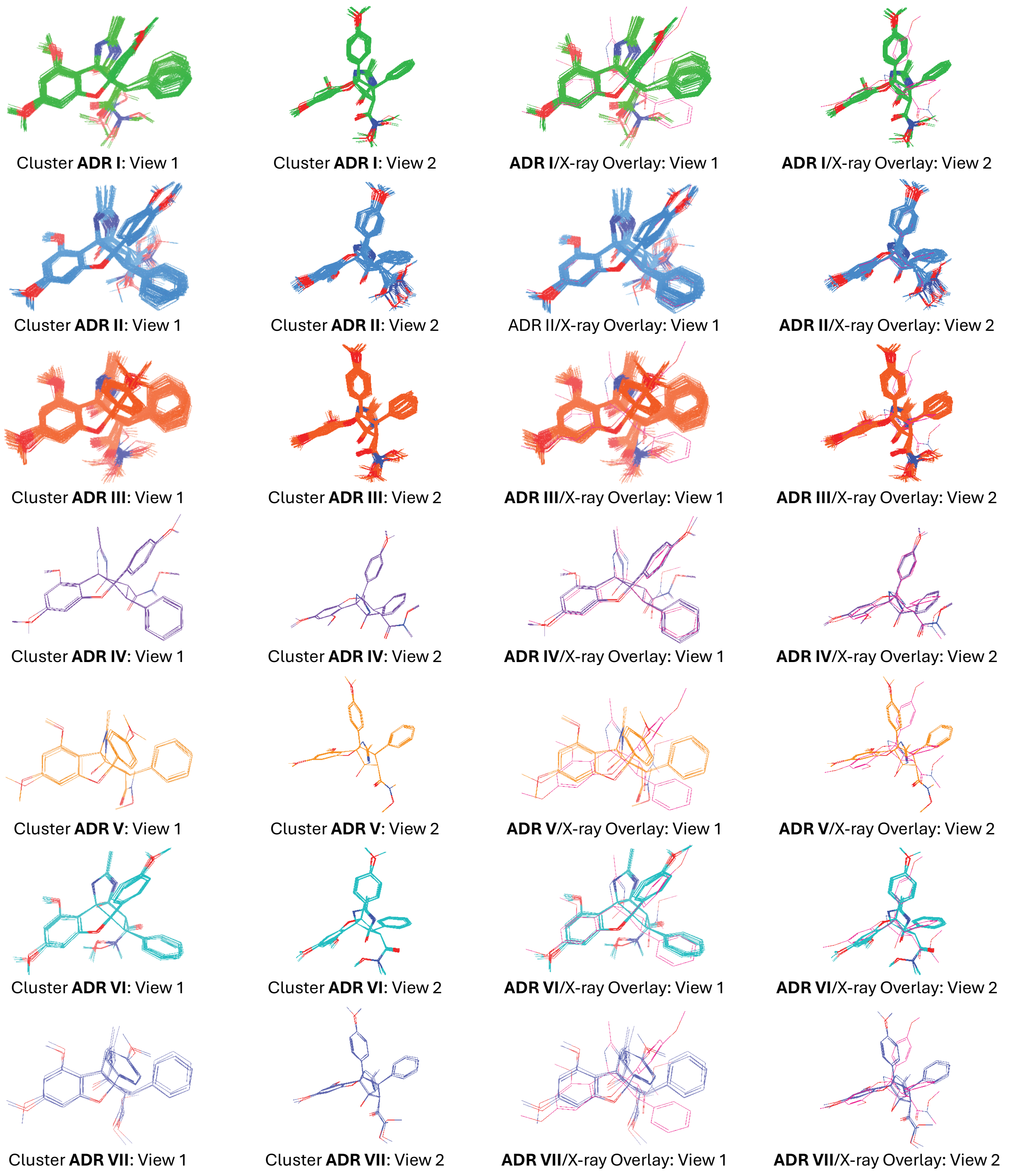


**Supplementary Figure S11.** DFT-optimized conformations for **CMLD012824**. Conformational clusters for **CMLD012824** after DFT (B3LYP-D3/6-31g**) geometry optimization. Overlays were generated *via* heavy atom superposition (Maestro version 2024-2, Schrödinger LLC) of each output conformer to the **CMLD012824** X-ray conformer (pink).

**Supplementary Table S1.** RocA conformational search results. Evaluation of force field quality and conformational search output for RocA reveals that the best-parameterized force fields (OPLS_2005 and OPLS4) fail to find the X-ray bound RocA conformers within a 5 kcal/mol energy window.

| Force Field | Stretch Parameters | | | Bend Parameters | | | Torsion Parameters | | | Lowest RMSD from X-ray (Å)[a] |
|---|---|---|---|---|---|---|---|---|---|---|
| | *High quality* | *Medium quality* | *Low quality* | *High quality* | *Medium quality* | *Low quality* | *High quality* | *Medium quality* | *Low quality* | |
| **MM3** | 20 | 1 | 14 | 40 | 13 | 11 | 28 | 29 | 21 | 0.23 |
| **MM2** | 62 | 4 | 0 | 121 | 11 | 0 | 57 | 79 | 9 | 0.22 |
| **AMBER** | 45 | 4 | 0 | 109 | 17 | 0 | 62 | 45 | 0 | 0.28 |
| **OPLS** | 54 | 0 | 0 | 116 | 4 | 6 | 69 | 36 | 6 | 0.21 |
| **MMFF** | 72 | 0 | 0 | 126 | 0 | 0 | 163 | 0 | 20 | 0.27 |
| **MMFFs** | 72 | 0 | 0 | 126 | 0 | 0 | 163 | 0 | 20 | 0.26 |
| **OPLS_2005** | 72 | 0 | 0 | 126 | 0 | 0 | 182 | 1 | 0 | 1.37 |
| **OPLS4** | 72 | 0 | 0 | 125 | 1 | 0 | 183 | 0 | 0 | 1.31 |

a. Lowest heavy atom root-mean-squared deviation (RMSD) from the X-ray conformation of RocA (PDB: 5ZC9) found among a conformer set generated *via* Macromodel conformational search (water solvent) with a 21.0 kJ/mol relative energy cutoff.

**Supplementary Table S2.** Summary of RocA conformational clusters

| Cluster | Cyclopentane configuration | Count (FF)[a] | Count (DFT)[b] | Computed properties for lowest-energy cluster member | | | |
|---|---|---|---|---|---|---|---|
| | | | | **Absolute energy (Hartrees)[c]** | **Relative energy (kcal/mol)[c]** | **Aryl-aryl torsion[d]** | **RMSD from X-ray (Å)** |
| *RocA X-ray* | *C3-exo$_{down}$* | - | - | - | - | *39.8°* | *0.00* |
| **RocA I** | C1-exo$_{up}$ | 16 | 12 | -1704.439766 | +2.37 | -12.1° | 1.73 |
| **RocA II** | C2-exo$_{down}$ | 4 | 4 | -1704.441238 | +1.45 | -34.2° | 2.29 |
| **RocA III** | C3-exo$_{down}$ | 4 | 4 | -1704.443552 | 0.00 | 41.2° | 0.59 |
| **RocA IV** | C8b-exo$_{down}$ | 0 | 4 | -1704.436322 | +4.53 | -21.8° | 1.19 |

a) count of cluster members following force field-based (FF) Macromodel conformational search; b) count of cluster members following density functional theory-based (DFT) geometry optimization; c) gas phase energies calculated using M06-2X/6-31g++**; d) Defined as the torsion angle between the rocaglate "B" and "C" aryl rings as previously reported.[S1]

**Supplementary Table S3.** Summary of **CMLD012824** conformational clusters

| Cluster | Cyclopentane configuration | Count (FF)[a] | Count (DFT)[b] | Computed properties for lowest-energy cluster member | | | |
|---|---|---|---|---|---|---|---|
| | | | | Absolute energy (Hartrees)[c] | Relative energy (kcal/mol)[c] | Aryl-aryl torsion[d] | RMSD from X-ray (Å) |
| *ADR X-ray* | *C3-exo$_{down}$* | - | - | - | - | *42.4°* | *0.00* |
| **ADR I** | C2-exo$_{down}$ | 44 | 49 | -1891.248342 | +3.87 | -11.5° | 1.77 |
| **ADR II** | C3-exo$_{down}$ | 43 | 48 | -1891.254516 | 0.00 | 42.0° | 0.55 |
| **ADR III** | C3-exo$_{up}$ | 76 | 69 | -1891.247400 | +4.47 | -30.5° | 2.19 |
| **ADR IV** | C2-exo$_{up}$ | 15 | 3 | -1891.248576 | +3.72 | 41.3° | 0.70 |
| **ADR V** | C8b-exo$_{up}$ | 2 | 3 | -1891.234632 | +12.48 | -26.0° | 2.15 |
| **ADR VI** | C3a-exo$_{up}$ | 1 | 9 | -1891.248988 | +3.47 | 32.3° | 1.58 |
| **ADR VII** | C3a-exo$_{down}$ | 0 | 3 | -1891.235825 | +11.73 | -32.2° | 2.05 |

a) count of cluster members following force field-based (FF) Macromodel conformational search; b) count of cluster members following density functional theory-based (DFT) geometry optimization; c) gas phase energies calculated using M06-2X/6-31g++**; d) Defined as the torsion angle between the rocaglate "B" and "C" aryl rings as previously reported.[S1]

**Computational Data**

Coordinates and gas phase energy (MO6-2X/6-31g++**) for the lowest-energy member of cluster **RocA I**

```
Gas phase energy......................   -1704.43976600690 hartrees

atom              x                  y                  z
  C3276        38.5655330000      7.5289190000      42.9514410000
  C3277        43.4260880000      7.6706260000      40.8124220000
  C3278        44.0931410000      8.8909940000      40.7312870000
  C3279        44.8291350000      9.3586680000      41.8381040000
  C3280        44.9303950000      8.6213240000      43.0194820000
  C3281        44.2416260000      7.4113400000      43.0513600000
  C3282        46.1406020000     11.1455910000      42.7257200000
  C3283        42.5335270000      7.8880260000      38.5923260000
  C3284        44.2673560000      4.1918850000      44.1140020000
  C3285        45.5040310000      4.2814280000      44.7538070000
  C3286        46.2620980000      3.1436460000      45.0458830000
  C3287        39.5880310000      9.6598360000      43.5681580000
  C3288        45.7787550000      1.8819670000      44.6914740000
  C3289        44.5427240000      1.7764590000      44.0392110000
  C3290        43.8033330000      2.9133720000      43.7562260000
  C3291        47.6695680000      0.7425850000      45.6047700000
  C3292        41.7824150000      4.3354450000      45.5930810000
  C3293        42.4818740000      4.0829980000      46.7815160000
  C3294        42.2438080000      2.9336110000      47.5299230000
  C3295        41.2871200000      2.0087200000      47.1085140000
  C3296        40.5740510000      2.2504400000      45.9364000000
  C3297        40.8218330000      3.4015510000      45.1871240000
  C3298        40.8852430000      7.6396920000      43.9495450000
  C3299        41.0003230000      6.1045820000      43.8857000000
  C3300        42.1377510000      5.5779590000      44.7972030000
  C3301        43.4203690000      5.4250120000      43.8701280000
  C3302        42.8899520000      5.5951930000      42.3718810000
  C3303        41.3515220000      5.5557190000      42.4836730000
  C3304        43.4889460000      6.9236530000      41.9948270000
  N3305        39.7087880000      8.2096300000      43.5367420000
  O3306        41.8050750000      8.3220950000      44.4004930000
  O3307        40.6615730000      6.2248980000      41.4533520000
  O3308        44.2579940000      6.5909020000      44.1285400000
  O3309        45.4235100000     10.5706120000      41.6425760000
  O3310        42.7023940000      7.1241720000      39.7812380000
  O3311        43.2292130000      4.5589060000      41.4590240000
  O3312        46.4233050000      0.7005460000      44.9288510000
  H38          38.4386830000      7.8255080000      41.9038800000
  H39          38.6829350000      6.4503130000      42.9617730000
  H40          37.6554930000      7.7948980000      43.5053370000
  H41          44.0701010000      9.5114420000      39.8461090000
  H42          45.4791520000      8.9557380000      43.8879240000
  H43          46.5038130000     12.1102870000      42.3676420000
  H44          45.4944900000     11.3014860000      43.5991050000
  H45          46.9964150000     10.5240440000      43.0200130000
  H46          41.9049120000      7.2848580000      37.9357760000
```

| | | | |
|---|---|---|---|
| H47 | 42.0381050000 | 8.8444620000 | 38.7988860000 |
| H48 | 43.4956830000 | 8.0759120000 | 38.1014410000 |
| H49 | 45.8829460000 | 5.2555660000 | 45.0384220000 |
| H50 | 47.2152550000 | 3.2582850000 | 45.5482170000 |
| H51 | 40.4821730000 | 10.0787360000 | 44.0253660000 |
| H52 | 39.4783880000 | 10.0568100000 | 42.5503580000 |
| H53 | 38.7044560000 | 9.9543600000 | 44.1485340000 |
| H54 | 44.1819350000 | 0.7898660000 | 43.7684860000 |
| H55 | 42.8507550000 | 2.8086890000 | 43.2539860000 |
| H56 | 47.9990350000 | -0.2939290000 | 45.6969100000 |
| H57 | 48.4211940000 | 1.3098740000 | 45.0393000000 |
| H58 | 47.5740040000 | 1.1807290000 | 46.6074210000 |
| H59 | 43.2350830000 | 4.7939230000 | 47.1099540000 |
| H60 | 42.8046960000 | 2.7593870000 | 48.4438290000 |
| H61 | 41.0990660000 | 1.1103340000 | 47.6893660000 |
| H62 | 39.8256430000 | 1.5397000000 | 45.5968870000 |
| H63 | 40.2633770000 | 3.5535960000 | 44.2685300000 |
| H64 | 40.0468990000 | 5.6858940000 | 44.2127830000 |
| H65 | 42.4090600000 | 6.3651650000 | 45.5022670000 |
| H66 | 41.0958300000 | 4.4827440000 | 42.4737590000 |
| H67 | 41.2327090000 | 6.2425700000 | 40.6687130000 |
| H68 | 44.1907800000 | 4.5648810000 | 41.3539220000 |

Coordinates and gas phase energy (MO6-2X/6-31g++**) for the lowest-energy member of cluster **RocA II**

Gas phase energy...................... -1704.44123833503 hartrees

| atom | x | y | z |
|---|---|---|---|
| C3276 | 39.8639180000 | 7.1828620000 | 46.4351450000 |
| C3277 | 43.8310720000 | 7.4196940000 | 40.6733560000 |
| C3278 | 44.5755530000 | 8.5995150000 | 40.6148140000 |
| C3279 | 45.0399230000 | 9.1856190000 | 41.8074930000 |
| C3280 | 44.7954070000 | 8.6118180000 | 43.0621250000 |
| C3281 | 44.0491260000 | 7.4380720000 | 43.0583380000 |
| C3282 | 46.2312650000 | 11.0156230000 | 42.7765920000 |
| C3283 | 43.4211030000 | 7.3600440000 | 38.3140820000 |
| C3284 | 44.1245560000 | 4.3791060000 | 44.1033970000 |
| C3285 | 45.2894580000 | 4.6711440000 | 44.8133870000 |
| C3286 | 46.2211660000 | 3.6793410000 | 45.1353470000 |
| C3287 | 40.5540040000 | 9.4745510000 | 45.9089140000 |
| C3288 | 45.9873020000 | 2.3601810000 | 44.7412770000 |
| C3289 | 44.8204790000 | 2.0514340000 | 44.0288880000 |
| C3290 | 43.9065850000 | 3.0447830000 | 43.7159940000 |
| C3291 | 48.0255050000 | 1.5631210000 | 45.6991890000 |
| C3292 | 41.2581580000 | 4.0253540000 | 44.9774340000 |
| C3293 | 40.3020600000 | 3.3339060000 | 44.2242660000 |
| C3294 | 39.8863570000 | 2.0539220000 | 44.5955160000 |
| C3295 | 40.4189690000 | 1.4414160000 | 45.7284850000 |
| C3296 | 41.3686240000 | 2.1214190000 | 46.4924200000 |
| C3297 | 41.7782410000 | 3.3980750000 | 46.1191320000 |
| C3298 | 40.8650630000 | 7.7657080000 | 44.1963140000 |
| C3299 | 40.7910660000 | 6.2873900000 | 43.7970170000 |

| | | | |
|---|---|---|---|
| C3300 | 41.7744030000 | 5.4097670000 | 44.6250170000 |
| C3301 | 43.1078720000 | 5.4647690000 | 43.8027580000 |
| C3302 | 42.7090770000 | 5.6692840000 | 42.2861590000 |
| C3303 | 41.1750380000 | 6.0163270000 | 42.3103060000 |
| C3304 | 43.5647530000 | 6.8429990000 | 41.9130670000 |
| N3305 | 40.4286820000 | 8.0965980000 | 45.4527580000 |
| O3306 | 41.2897360000 | 8.6328220000 | 43.4228240000 |
| O3307 | 40.7585740000 | 6.9829870000 | 41.3843020000 |
| O3308 | 43.6840110000 | 6.7633690000 | 44.1902170000 |
| O3309 | 45.7505720000 | 10.3383030000 | 41.6265080000 |
| O3310 | 43.3441280000 | 6.7405430000 | 39.5904950000 |
| O3311 | 42.9408180000 | 4.5148310000 | 41.4949000000 |
| O3312 | 46.8203750000 | 1.3063050000 | 44.9977660000 |
| H38 | 40.5399440000 | 7.0634760000 | 47.2924600000 |
| H39 | 38.9146730000 | 7.5868120000 | 46.8078190000 |
| H40 | 39.6701180000 | 6.1968240000 | 46.0193770000 |
| H41 | 44.8195430000 | 9.0900530000 | 39.6821690000 |
| H42 | 45.1419840000 | 9.0415280000 | 43.9911530000 |
| H43 | 46.7442240000 | 11.9063710000 | 42.4096450000 |
| H44 | 45.4108490000 | 11.3181850000 | 43.4407690000 |
| H45 | 46.9409700000 | 10.3989560000 | 43.3442400000 |
| H46 | 42.9036270000 | 6.6932160000 | 37.6228350000 |
| H47 | 42.9235220000 | 8.3372130000 | 38.3191930000 |
| H48 | 44.4617530000 | 7.4839780000 | 37.9886410000 |
| H49 | 45.4791490000 | 5.6916150000 | 45.1237460000 |
| H50 | 47.1139700000 | 3.9513580000 | 45.6859090000 |
| H51 | 41.0209220000 | 10.0652440000 | 45.1234120000 |
| H52 | 39.5665660000 | 9.8918480000 | 46.1421090000 |
| H53 | 41.1700550000 | 9.5162220000 | 46.8159150000 |
| H54 | 44.6528300000 | 1.0217970000 | 43.7307920000 |
| H55 | 43.0137840000 | 2.7854450000 | 43.1633500000 |
| H56 | 48.5361340000 | 0.6022400000 | 45.7866710000 |
| H57 | 48.6714920000 | 2.2657960000 | 45.1555640000 |
| H58 | 47.8353030000 | 1.9617640000 | 46.7051430000 |
| H59 | 39.8745110000 | 3.7814220000 | 43.3340810000 |
| H60 | 39.1457640000 | 1.5357330000 | 43.9929960000 |
| H61 | 40.0972510000 | 0.4442080000 | 46.0149010000 |
| H62 | 41.7937350000 | 1.6557940000 | 47.3769650000 |
| H63 | 42.5326320000 | 3.9141240000 | 46.7066120000 |
| H64 | 39.7587750000 | 5.9619640000 | 43.9563660000 |
| H65 | 42.0227500000 | 5.8989280000 | 45.5694680000 |
| H66 | 40.6897280000 | 5.0827040000 | 42.0106900000 |
| H67 | 41.0497000000 | 7.8349740000 | 41.7622240000 |
| H68 | 42.9607290000 | 4.8274780000 | 40.5779740000 |

Coordinates and gas phase energy (MO6-2X/6-31g++**) for the lowest-energy member of cluster **RocA III**

Gas phase energy...................... -1704.44355206986 hartrees

| atom | x | y | z |
|---|---|---|---|
| C3276 | 37.4654770000 | 6.4619650000 | 43.5747250000 |
| C3277 | 43.7984050000 | 8.2884000000 | 41.0014140000 |

| | | | |
|---|---|---|---|
| C3278 | 44.5954160000 | 9.4200590000 | 41.1564340000 |
| C3279 | 45.4291260000 | 9.5401570000 | 42.2863170000 |
| C3280 | 45.4852700000 | 8.5455150000 | 43.2655210000 |
| C3281 | 44.6711890000 | 7.4313750000 | 43.0620200000 |
| C3282 | 47.0061370000 | 10.9117320000 | 43.4439940000 |
| C3283 | 42.8974660000 | 9.0752520000 | 38.9160650000 |
| C3284 | 44.0824650000 | 4.0544520000 | 43.6572660000 |
| C3285 | 45.0651450000 | 3.7678060000 | 44.6088610000 |
| C3286 | 45.4611860000 | 2.4584240000 | 44.8822270000 |
| C3287 | 38.6323940000 | 4.9324480000 | 42.0400860000 |
| C3288 | 44.8597760000 | 1.3965380000 | 44.1991980000 |
| C3289 | 43.8762590000 | 1.6672120000 | 43.2391900000 |
| C3290 | 43.4966440000 | 2.9764500000 | 42.9733950000 |
| C3291 | 46.1273140000 | -0.2579510000 | 45.3674460000 |
| C3292 | 42.3038540000 | 5.2636920000 | 45.7215670000 |
| C3293 | 41.9308200000 | 3.9441600000 | 46.0053390000 |
| C3294 | 41.9903760000 | 3.4491390000 | 47.3066010000 |
| C3295 | 42.4290460000 | 4.2654550000 | 48.3492240000 |
| C3296 | 42.8062870000 | 5.5809760000 | 48.0787970000 |
| C3297 | 42.7417000000 | 6.0724740000 | 46.7760950000 |
| C3298 | 39.8513180000 | 6.1286740000 | 43.8519900000 |
| C3299 | 41.1487290000 | 5.4389850000 | 43.3953180000 |
| C3300 | 42.3134800000 | 5.8187030000 | 44.3180420000 |
| C3301 | 43.5876340000 | 5.4724940000 | 43.4877100000 |
| C3302 | 43.1761530000 | 5.9176040000 | 42.0209980000 |
| C3303 | 41.6128990000 | 5.8606350000 | 41.9729800000 |
| C3304 | 43.8242260000 | 7.2788730000 | 41.9721080000 |
| N3305 | 38.6780570000 | 5.7156260000 | 43.2649160000 |
| O3306 | 39.8745690000 | 6.9784240000 | 44.7357250000 |
| O3307 | 40.9819740000 | 7.0851490000 | 41.6523430000 |
| O3308 | 44.6522370000 | 6.3750780000 | 43.9156330000 |
| O3309 | 46.1551100000 | 10.6928480000 | 42.3271840000 |
| O3310 | 42.9570460000 | 8.0825200000 | 39.9368690000 |
| O3311 | 43.6320210000 | 5.0747130000 | 40.9641140000 |
| O3312 | 45.1557960000 | 0.0769400000 | 44.3882870000 |
| H38 | 37.5966910000 | 6.9603940000 | 44.5333350000 |
| H39 | 37.2592110000 | 7.2218920000 | 42.8079700000 |
| H40 | 36.6128090000 | 5.7765380000 | 43.6267370000 |
| H41 | 44.5993070000 | 10.2331280000 | 40.4435950000 |
| H42 | 46.1145220000 | 8.6053060000 | 44.1422250000 |
| H43 | 47.4720300000 | 11.8847220000 | 43.2804360000 |
| H44 | 46.4403170000 | 10.9340310000 | 44.3842230000 |
| H45 | 47.7872800000 | 10.1435750000 | 43.5135160000 |
| H46 | 42.1859620000 | 8.7022620000 | 38.1782710000 |
| H47 | 42.5466280000 | 10.0358510000 | 39.3120200000 |
| H48 | 43.8764870000 | 9.2118740000 | 38.4426330000 |
| H49 | 45.5181650000 | 4.5815800000 | 45.1621940000 |
| H50 | 46.2220190000 | 2.2829380000 | 45.6331040000 |
| H51 | 38.8067390000 | 5.5574490000 | 41.1543310000 |
| H52 | 39.3725090000 | 4.1307700000 | 42.0523360000 |
| H53 | 37.6471240000 | 4.4652140000 | 41.9549910000 |
| H54 | 43.4266510000 | 0.8337460000 | 42.7097120000 |
| H55 | 42.7498890000 | 3.1599770000 | 42.2108320000 |

| | | | |
|---|---|---|---|
| H56 | 46.1988080000 | -1.3469880000 | 45.3615620000 |
| H57 | 47.1105440000 | 0.1684860000 | 45.1274910000 |
| H58 | 45.8276260000 | 0.0787690000 | 46.3687830000 |
| H59 | 41.6137820000 | 3.2845860000 | 45.2040930000 |
| H60 | 41.6977330000 | 2.4216210000 | 47.5044830000 |
| H61 | 42.4747750000 | 3.8800250000 | 49.3640730000 |
| H62 | 43.1460830000 | 6.2271470000 | 48.8836210000 |
| H63 | 43.0313720000 | 7.0984740000 | 46.5677660000 |
| H64 | 41.0022610000 | 4.3542440000 | 43.4052540000 |
| H65 | 42.2929140000 | 6.9107490000 | 44.3883860000 |
| H66 | 41.3602320000 | 5.0917750000 | 41.2306130000 |
| H67 | 41.4068720000 | 7.4336950000 | 40.8519540000 |
| H68 | 44.5943330000 | 5.0083450000 | 41.0395860000 |

Coordinates and gas phase energy (MO6-2X/6-31g++**) for the lowest-energy member of cluster **RocA IV**

Gas phase energy............. -1704.43632201431 hartrees

| atom | x | y | z |
|---|---|---|---|
| C3276 | 38.5952380000 | 8.5076530000 | 43.0257580000 |
| C3277 | 43.2363990000 | 8.1673440000 | 41.1735720000 |
| C3278 | 43.8783080000 | 9.3954490000 | 41.3087810000 |
| C3279 | 44.8816020000 | 9.5575060000 | 42.2812450000 |
| C3280 | 45.2464850000 | 8.5126520000 | 43.1334460000 |
| C3281 | 44.5721620000 | 7.3075620000 | 42.9561190000 |
| C3282 | 46.4296800000 | 11.0657320000 | 43.2969630000 |
| C3283 | 41.6732760000 | 9.0599910000 | 39.5715140000 |
| C3284 | 44.2626030000 | 3.8926830000 | 43.7414270000 |
| C3285 | 45.0776880000 | 3.7167300000 | 44.8650410000 |
| C3286 | 45.5387220000 | 2.4598740000 | 45.2499680000 |
| C3287 | 38.4322060000 | 6.0527770000 | 42.7432980000 |
| C3288 | 45.1782830000 | 1.3352550000 | 44.5000190000 |
| C3289 | 44.3774690000 | 1.4953380000 | 43.3642320000 |
| C3290 | 43.9299240000 | 2.7570400000 | 42.9902640000 |
| C3291 | 46.3629430000 | -0.1668600000 | 45.9311450000 |
| C3292 | 42.3179490000 | 4.9440970000 | 45.6687540000 |
| C3293 | 42.7139590000 | 5.5251470000 | 46.8779870000 |
| C3294 | 42.6195350000 | 4.8198950000 | 48.0786660000 |
| C3295 | 42.1249280000 | 3.5167210000 | 48.0860540000 |
| C3296 | 41.7232960000 | 2.9258110000 | 46.8861960000 |
| C3297 | 41.8175720000 | 3.6346280000 | 45.6918330000 |
| C3298 | 40.5751620000 | 7.1487590000 | 43.4319170000 |
| C3299 | 41.2052210000 | 5.7439810000 | 43.4844010000 |
| C3300 | 42.4745910000 | 5.7171690000 | 44.3824590000 |
| C3301 | 43.7172420000 | 5.2759070000 | 43.4686740000 |
| C3302 | 43.1969650000 | 5.6231260000 | 42.0384510000 |
| C3303 | 41.7075490000 | 5.2506010000 | 42.0997130000 |
| C3304 | 43.5911760000 | 7.0941310000 | 41.9940280000 |
| N3305 | 39.2725480000 | 7.2176250000 | 43.0054210000 |
| O3306 | 41.2097280000 | 8.1575470000 | 43.7373750000 |
| O3307 | 40.9748160000 | 5.6073650000 | 40.9480780000 |
| O3308 | 44.7945320000 | 6.2224080000 | 43.7452590000 |

| | | | |
|---|---|---|---|
| O3309 | 45.4321830000 | 10.8045910000 | 42.3202070000 |
| O3310 | 42.2243550000 | 7.9433800000 | 40.2650770000 |
| O3311 | 43.8312680000 | 4.9006940000 | 40.9873260000 |
| O3312 | 45.5554410000 | 0.0526900000 | 44.7853330000 |
| H38 | 39.3272930000 | 9.2827110000 | 43.2452810000 |
| H39 | 38.1285490000 | 8.7059300000 | 42.0537630000 |
| H40 | 37.8117910000 | 8.5264580000 | 43.7947550000 |
| H41 | 43.6251250000 | 10.2544400000 | 40.7031600000 |
| H42 | 45.9880110000 | 8.6090830000 | 43.9137630000 |
| H43 | 46.7254140000 | 12.1064130000 | 43.1542780000 |
| H44 | 46.0411050000 | 10.9362600000 | 44.3154750000 |
| H45 | 47.3064140000 | 10.4187070000 | 43.1632090000 |
| H46 | 41.3072640000 | 9.8166540000 | 40.2748140000 |
| H47 | 42.4099690000 | 9.5103820000 | 38.8969680000 |
| H48 | 40.8414160000 | 8.6665940000 | 38.9856720000 |
| H49 | 45.3445860000 | 4.5800740000 | 45.4632990000 |
| H50 | 46.1613730000 | 2.3716690000 | 46.1319490000 |
| H51 | 38.9612510000 | 5.3178320000 | 42.1385100000 |
| H52 | 38.0671340000 | 5.5891960000 | 43.6702540000 |
| H53 | 37.5620330000 | 6.3799190000 | 42.1684220000 |
| H54 | 44.1245540000 | 0.6160940000 | 42.7810570000 |
| H55 | 43.3467830000 | 2.8559970000 | 42.0850490000 |
| H56 | 46.5396510000 | -1.2430880000 | 45.9751780000 |
| H57 | 47.3275600000 | 0.3529240000 | 45.8557200000 |
| H58 | 45.8579940000 | 0.1534020000 | 46.8525160000 |
| H59 | 43.1043800000 | 6.5394510000 | 46.8742630000 |
| H60 | 42.9322360000 | 5.2901350000 | 49.0069580000 |
| H61 | 42.0526390000 | 2.9638730000 | 49.0184640000 |
| H62 | 41.3449750000 | 1.9074790000 | 46.8802390000 |
| H63 | 41.5352970000 | 3.1508160000 | 44.7610360000 |
| H64 | 40.4577400000 | 5.0446070000 | 43.8628180000 |
| H65 | 42.6829560000 | 6.7575380000 | 44.6368370000 |
| H66 | 41.6735610000 | 4.1567260000 | 42.1239590000 |
| H67 | 41.1637160000 | 6.5434700000 | 40.7514140000 |
| H68 | 43.2575180000 | 5.0223640000 | 40.2150830000 |

Coordinates and gas phase energy (MO6-2X/6-31g++**) for the lowest-energy member of cluster **ADR I**

Gas phase energy...................... -1891.24834231225 hartrees

| atom | x | y | z |
|---|---|---|---|
| C1 | -28.4813740000 | 18.5948020000 | 44.0484690000 |
| C2 | -33.9814450000 | 22.0554170000 | 40.0959420000 |
| C3 | -32.6613600000 | 22.4118860000 | 40.3607660000 |
| C4 | -30.4154620000 | 23.8997530000 | 42.3631630000 |
| C5 | -31.6489470000 | 24.7938010000 | 42.3775540000 |
| C6 | -32.8216820000 | 24.4277390000 | 43.0655340000 |
| C7 | -33.9306890000 | 25.2586030000 | 43.1031210000 |
| C8 | -33.9073150000 | 26.4972430000 | 42.4477720000 |
| C9 | -35.0902790000 | 28.4941780000 | 41.8825890000 |
| C10 | -32.7544270000 | 26.8827120000 | 41.7602060000 |
| C11 | -31.6438120000 | 26.0338060000 | 41.7352050000 |

| | | | |
|---|---|---|---|
| C12 | -26.7411550000 | 19.4139980000 | 41.4924240000 |
| C13 | -28.3334660000 | 24.8880950000 | 42.2761040000 |
| C14 | -27.2399700000 | 25.5774210000 | 41.7585870000 |
| C15 | -26.1927710000 | 25.8304810000 | 42.6487230000 |
| C16 | -24.8824770000 | 26.8914570000 | 40.9568220000 |
| C17 | -26.2423810000 | 25.4124780000 | 43.9928240000 |
| C18 | -27.3588610000 | 24.7202890000 | 44.4588320000 |
| C19 | -26.4104060000 | 24.4320500000 | 46.6444080000 |
| C20 | -28.4284350000 | 24.4745490000 | 43.5933120000 |
| C21 | -29.7061900000 | 23.7259570000 | 43.7880650000 |
| C22 | -31.1080240000 | 22.9629890000 | 45.4265670000 |
| C23 | -28.3923630000 | 21.1851650000 | 42.0158620000 |
| C24 | -32.1892410000 | 23.0823410000 | 46.4539200000 |
| C25 | -29.5764320000 | 22.1703860000 | 44.0292570000 |
| C26 | -29.7492210000 | 21.4813700000 | 42.6570280000 |
| C27 | -30.5855200000 | 22.4370250000 | 41.7684550000 |
| C28 | -32.0112410000 | 21.9762440000 | 41.5230610000 |
| C29 | -32.7208100000 | 21.1597110000 | 42.4141540000 |
| C30 | -34.0431910000 | 20.7999510000 | 42.1503740000 |
| C31 | -34.6797110000 | 21.2457280000 | 40.9927280000 |
| N32 | -27.8917500000 | 19.9193930000 | 42.2124910000 |
| N33 | -30.4853530000 | 24.1052720000 | 44.9548010000 |
| N34 | -30.6745150000 | 21.8440060000 | 44.9542040000 |
| O35 | -28.7458140000 | 18.9012760000 | 42.6672590000 |
| O36 | -27.7591920000 | 22.0169300000 | 41.3737530000 |
| O37 | -35.0498100000 | 27.2387690000 | 42.5434490000 |
| O38 | -29.4183240000 | 24.5610650000 | 41.5288060000 |
| O39 | -25.0461810000 | 26.4883220000 | 42.3084900000 |
| O40 | -27.4901400000 | 24.2357880000 | 45.7381640000 |
| O41 | -28.3452630000 | 21.7775520000 | 44.5897570000 |
| H42 | -32.2548560000 | 20.8342660000 | 43.3386530000 |
| H43 | -34.5771680000 | 20.1705330000 | 42.8572170000 |
| H44 | -34.4646630000 | 22.4088970000 | 39.1893250000 |
| H45 | -35.7090630000 | 20.9640010000 | 40.7892780000 |
| H46 | -32.1254050000 | 23.0517690000 | 39.6643870000 |
| H47 | -30.7581890000 | 26.3443100000 | 41.1953170000 |
| H48 | -32.7030010000 | 27.8322060000 | 41.2407470000 |
| H49 | -32.8843430000 | 23.4650690000 | 43.5539470000 |
| H50 | -34.8355310000 | 24.9622650000 | 43.6234240000 |
| H51 | -27.2206060000 | 25.8635760000 | 40.7168360000 |
| H52 | -25.3883220000 | 25.6328330000 | 44.6181920000 |
| H53 | -29.1576070000 | 17.7695070000 | 44.2842950000 |
| H54 | -27.4431920000 | 18.2668660000 | 44.1838300000 |
| H55 | -28.6831790000 | 19.4582790000 | 44.6861490000 |
| H56 | -36.0809100000 | 28.9075560000 | 42.0796910000 |
| H57 | -34.3285500000 | 29.1832370000 | 42.2714430000 |
| H58 | -34.9534130000 | 28.3869590000 | 40.7983200000 |
| H59 | -26.1613340000 | 20.2708300000 | 41.1517280000 |
| H60 | -26.1307590000 | 18.7949800000 | 42.1565400000 |
| H61 | -27.0557710000 | 18.8128720000 | 40.6310650000 |
| H62 | -23.9026890000 | 27.3689640000 | 40.9013940000 |
| H63 | -24.9070510000 | 26.0328340000 | 40.2734490000 |
| H64 | -25.6528750000 | 27.6124610000 | 40.6527660000 |

| | | | |
|---|---|---|---|
| H65 | -26.7173230000 | 23.9657890000 | 47.5819910000 |
| H66 | -25.4909180000 | 23.9559860000 | 46.2820830000 |
| H67 | -26.2227920000 | 25.4990730000 | 46.8137010000 |
| H68 | -31.8056290000 | 23.5725010000 | 47.3551870000 |
| H69 | -33.0134900000 | 23.6942090000 | 46.0702950000 |
| H70 | -32.5628380000 | 22.0911020000 | 46.7114770000 |
| H71 | -30.2646450000 | 20.5374050000 | 42.8115650000 |
| H72 | -30.0992720000 | 22.5103570000 | 40.7936590000 |
| H73 | -30.9825970000 | 24.9856040000 | 44.9457170000 |
| H74 | -28.1505360000 | 22.3792540000 | 45.3266730000 |

Coordinates and gas phase energy (MO6-2X/6-31g++**) for the lowest-energy member of cluster **ADR II**

Gas phase energy...................... -1891.25451617524 hartrees

| atom | x | y | z |
|---|---|---|---|
| C1 | -31.8957830000 | 19.6951690000 | 45.0727020000 |
| C2 | -33.0486720000 | 21.6374240000 | 39.8448630000 |
| C3 | -32.1834930000 | 21.8008010000 | 40.9271240000 |
| C4 | -30.1756250000 | 24.1564360000 | 42.5933930000 |
| C5 | -31.6205310000 | 24.5726920000 | 42.7869340000 |
| C6 | -32.4279490000 | 24.0581990000 | 43.8064180000 |
| C7 | -33.7897730000 | 24.3598340000 | 43.8872310000 |
| C8 | -34.3715890000 | 25.1979490000 | 42.9327850000 |
| C9 | -36.5524380000 | 25.0149510000 | 43.8908210000 |
| C10 | -33.5757660000 | 25.7282410000 | 41.9105690000 |
| C11 | -32.2266890000 | 25.4156170000 | 41.8408010000 |
| C12 | -30.6255140000 | 18.0403940000 | 42.6430080000 |
| C13 | -28.2918640000 | 25.3991990000 | 42.2219210000 |
| C14 | -27.3571450000 | 26.1908280000 | 41.5552140000 |
| C15 | -26.1342600000 | 26.3806610000 | 42.2092050000 |
| C16 | -25.2860750000 | 27.7391120000 | 40.4345980000 |
| C17 | -25.8666560000 | 25.8120920000 | 43.4696900000 |
| C18 | -26.8365080000 | 25.0288930000 | 44.0971150000 |
| C19 | -25.4152370000 | 24.4252010000 | 45.9163680000 |
| C20 | -28.0592320000 | 24.8099660000 | 43.4525180000 |
| C21 | -29.2471230000 | 24.0113970000 | 43.8764060000 |
| C22 | -29.7203040000 | 23.2956670000 | 46.0191460000 |
| C23 | -29.7647130000 | 20.3688090000 | 42.7701720000 |
| C24 | -30.1077580000 | 23.4690790000 | 47.4545180000 |
| C25 | -29.0358700000 | 22.4385240000 | 44.0936790000 |
| C26 | -30.0774680000 | 21.8026950000 | 43.1246730000 |
| C27 | -30.0286820000 | 22.7484960000 | 41.9164970000 |
| C28 | -30.9773000000 | 22.4939230000 | 40.7727100000 |
| C29 | -30.6593540000 | 23.0189780000 | 39.5132850000 |
| C30 | -31.5253050000 | 22.8609090000 | 38.4337720000 |
| C31 | -32.7251330000 | 22.1668540000 | 38.5963140000 |
| N32 | -30.6769810000 | 19.4115950000 | 43.1078050000 |
| N33 | -29.7897430000 | 24.3915280000 | 45.1724500000 |
| N34 | -29.3815160000 | 22.1722680000 | 45.4904230000 |
| O35 | -31.9194390000 | 19.7863440000 | 43.6328490000 |
| O36 | -28.6983150000 | 20.0586170000 | 42.2222560000 |

| | | | |
|---|---|---|---|
| O37 | -35.6895840000 | 25.5583610000 | 42.9056100000 |
| O38 | -29.5302570000 | 25.1280790000 | 41.7209250000 |
| O39 | -25.1088870000 | 27.1258950000 | 41.7024150000 |
| O40 | -26.7074620000 | 24.4673150000 | 45.3286740000 |
| O41 | -27.7230060000 | 22.0670090000 | 43.8051440000 |
| H42 | -29.7294310000 | 23.5669190000 | 39.3914630000 |
| H43 | -31.2620050000 | 23.2758610000 | 37.4647070000 |
| H44 | -33.9802960000 | 21.0951730000 | 39.9810630000 |
| H45 | -33.4008160000 | 22.0386020000 | 37.7553240000 |
| H46 | -32.4562030000 | 21.3866340000 | 41.8913480000 |
| H47 | -31.6287010000 | 25.8182870000 | 41.0330590000 |
| H48 | -34.0413910000 | 26.3742190000 | 41.1736600000 |
| H49 | -32.0080850000 | 23.4132130000 | 44.5661370000 |
| H50 | -34.3755640000 | 23.9369050000 | 44.6946130000 |
| H51 | -27.5912330000 | 26.6199740000 | 40.5913160000 |
| H52 | -24.9019680000 | 26.0098410000 | 43.9170810000 |
| H53 | -32.8937150000 | 20.0083310000 | 45.3883150000 |
| H54 | -31.7162940000 | 18.6623860000 | 45.3934560000 |
| H55 | -31.1309130000 | 20.3590790000 | 45.4905890000 |
| H56 | -37.5467880000 | 25.4087750000 | 43.6728730000 |
| H57 | -36.5815600000 | 23.9178270000 | 43.8453360000 |
| H58 | -36.2590310000 | 25.3211980000 | 44.9041530000 |
| H59 | -29.6375820000 | 17.8744780000 | 42.2150830000 |
| H60 | -30.7859910000 | 17.3529060000 | 43.4786530000 |
| H61 | -31.3965210000 | 17.8740900000 | 41.8833840000 |
| H62 | -24.3572120000 | 28.2726750000 | 40.2256780000 |
| H63 | -25.4610370000 | 26.9956610000 | 39.6459080000 |
| H64 | -26.1194090000 | 28.4540770000 | 40.4428890000 |
| H65 | -25.0546310000 | 25.4289530000 | 46.1778590000 |
| H66 | -25.5189910000 | 23.8313630000 | 46.8256350000 |
| H67 | -24.6913460000 | 23.9439820000 | 45.2479870000 |
| H68 | -29.4638060000 | 24.2151740000 | 47.9343060000 |
| H69 | -31.1406570000 | 23.8249940000 | 47.5347010000 |
| H70 | -30.0050320000 | 22.5202870000 | 47.9814840000 |
| H71 | -31.0536600000 | 21.8631860000 | 43.5933680000 |
| H72 | -29.0030230000 | 22.7150380000 | 41.5332210000 |
| H73 | -29.5680870000 | 25.3096950000 | 45.5324230000 |
| H74 | -27.7410540000 | 21.2880390000 | 43.2159440000 |

Coordinates and gas phase energy (MO6-2X/6-31g++**) for the lowest-energy member of cluster **ADR III**

Gas phase energy...................... -1891.24740039854 hartrees

| atom | x | y | z |
|---|---|---|---|
| C1 | -30.9902060000 | 18.8478160000 | 40.6001980000 |
| C2 | -34.6994310000 | 22.3159200000 | 41.4272820000 |
| C3 | -33.3498930000 | 22.6097750000 | 41.2534630000 |
| C4 | -30.4466810000 | 23.9239800000 | 42.3966560000 |
| C5 | -31.4826390000 | 25.0366540000 | 42.4229410000 |
| C6 | -32.5924270000 | 24.9901270000 | 43.2872830000 |
| C7 | -33.5326230000 | 26.0087480000 | 43.3052050000 |
| C8 | -33.3972430000 | 27.1128310000 | 42.4532720000 |

| | | | |
|---|---|---|---|
| C9 | -34.3048640000 | 29.1887920000 | 41.6960430000 |
| C10 | -32.3052760000 | 27.1769540000 | 41.5850150000 |
| C11 | -31.3645140000 | 26.1430870000 | 41.5801150000 |
| C12 | -28.4914900000 | 20.2597690000 | 39.1670280000 |
| C13 | -28.3263530000 | 24.7881840000 | 42.1439040000 |
| C14 | -27.2433560000 | 25.4310520000 | 41.5540430000 |
| C15 | -26.2260240000 | 25.8413290000 | 42.4256870000 |
| C16 | -24.9234080000 | 26.7315540000 | 40.6313590000 |
| C17 | -26.3000900000 | 25.6222530000 | 43.8133890000 |
| C18 | -27.4075550000 | 24.9639720000 | 44.3545060000 |
| C19 | -26.4993930000 | 24.9243050000 | 46.5637060000 |
| C20 | -28.4320670000 | 24.5501720000 | 43.5020460000 |
| C21 | -29.6368830000 | 23.7021270000 | 43.7323330000 |
| C22 | -30.5395860000 | 22.6858650000 | 45.5796970000 |
| C23 | -28.8265600000 | 21.2266670000 | 41.4212700000 |
| C24 | -31.3003090000 | 22.6178440000 | 46.8665710000 |
| C25 | -29.3268150000 | 22.1312130000 | 43.8014020000 |
| C26 | -29.8861180000 | 21.5185760000 | 42.4769530000 |
| C27 | -30.9320390000 | 22.5209350000 | 41.9007020000 |
| C28 | -32.3828630000 | 22.1494500000 | 42.1580340000 |
| C29 | -32.8061720000 | 21.3700640000 | 43.2416380000 |
| C30 | -34.1595840000 | 21.0730540000 | 43.4175120000 |
| C31 | -35.1116280000 | 21.5447110000 | 42.5150740000 |
| N32 | -29.2218820000 | 20.3670640000 | 40.4178400000 |
| N33 | -30.3787050000 | 23.9141210000 | 44.9593930000 |
| N34 | -30.0967570000 | 21.6539120000 | 44.9529430000 |
| O35 | -30.6081390000 | 20.1833230000 | 40.2529160000 |
| O36 | -27.6765280000 | 21.6694030000 | 41.4292540000 |
| O37 | -34.3811140000 | 28.0574720000 | 42.5478740000 |
| O38 | -29.3920340000 | 24.2946210000 | 41.4411950000 |
| O39 | -25.0947660000 | 26.4897370000 | 42.0190780000 |
| O40 | -27.5920890000 | 24.7078610000 | 45.6819720000 |
| O41 | -27.9927590000 | 21.8201610000 | 44.0713760000 |
| H42 | -32.0833530000 | 21.0068450000 | 43.9643810000 |
| H43 | -34.4671870000 | 20.4687900000 | 44.2668430000 |
| H44 | -35.4288710000 | 22.6893880000 | 40.7141180000 |
| H45 | -36.1636500000 | 21.3119990000 | 42.6547280000 |
| H46 | -33.0369190000 | 23.2198730000 | 40.4105970000 |
| H47 | -30.5241890000 | 26.2030550000 | 40.8994580000 |
| H48 | -32.1716580000 | 28.0155940000 | 40.9121000000 |
| H49 | -32.7293200000 | 24.1412990000 | 43.9424060000 |
| H50 | -34.3916640000 | 25.9666570000 | 43.9668040000 |
| H51 | -27.2089460000 | 25.5810100000 | 40.4844740000 |
| H52 | -25.4810990000 | 25.9784660000 | 44.4234480000 |
| H53 | -32.0715620000 | 18.8182120000 | 40.4523670000 |
| H54 | -30.5040840000 | 18.1111900000 | 39.9495180000 |
| H55 | -30.7538610000 | 18.6241610000 | 41.6459870000 |
| H56 | -34.3362160000 | 28.9030230000 | 40.6358990000 |
| H57 | -35.1781930000 | 29.8014360000 | 41.9272910000 |
| H58 | -33.3947790000 | 29.7760010000 | 41.8786990000 |
| H59 | -28.8638180000 | 20.9888250000 | 38.4384910000 |
| H60 | -27.4404560000 | 20.4531380000 | 39.3767810000 |
| H61 | -28.6091330000 | 19.2511330000 | 38.7630780000 |

| | | | |
|---|---|---|---|
| H62 | -23.9632090000 | 27.2404430000 | 40.5302830000 |
| H63 | -24.8994580000 | 25.7961480000 | 40.0570560000 |
| H64 | -25.7171440000 | 27.3760990000 | 40.2305970000 |
| H65 | -26.8260550000 | 24.5659430000 | 47.5413080000 |
| H66 | -25.6149560000 | 24.3588710000 | 46.2464700000 |
| H67 | -26.2412240000 | 25.9889070000 | 46.6367130000 |
| H68 | -31.2632830000 | 21.6030150000 | 47.2630410000 |
| H69 | -30.8772360000 | 23.3091490000 | 47.6040200000 |
| H70 | -32.3441980000 | 22.9079370000 | 46.7030750000 |
| H71 | -30.3668300000 | 20.5729700000 | 42.7263710000 |
| H72 | -30.8315650000 | 22.5698990000 | 40.8149770000 |
| H73 | -30.1565200000 | 24.7261650000 | 45.5189810000 |
| H74 | -27.5091340000 | 21.9306890000 | 43.2300880000 |

Coordinates and gas phase energy (MO6-2X/6-31g++**) for the lowest-energy member of cluster **ADR IV**

Gas phase energy...................... -1891.24857560840 hartrees

| atom | x | y | z |
|---|---|---|---|
| C3233 | -29.7095950000 | 20.3366620000 | 42.8332640000 |
| C3234 | -30.2317000000 | 17.9461980000 | 43.1785890000 |
| C3235 | -32.9380910000 | 19.4232140000 | 43.2437820000 |
| C3236 | -30.0490320000 | 21.7670200000 | 43.1842370000 |
| C3237 | -30.0827690000 | 22.7190530000 | 41.9726840000 |
| C3238 | -31.0544050000 | 22.3909900000 | 40.8663760000 |
| C3239 | -30.5709840000 | 22.2535280000 | 39.5599680000 |
| C3240 | -31.4261190000 | 21.9422030000 | 38.5029120000 |
| C3241 | -32.7884060000 | 21.7621080000 | 38.7366280000 |
| C3242 | -33.2854190000 | 21.8974670000 | 40.0339440000 |
| C3243 | -32.4276170000 | 22.2077790000 | 41.0879960000 |
| C3244 | -28.9475200000 | 22.4295740000 | 44.0798020000 |
| C3245 | -29.2262610000 | 23.9933850000 | 43.9000970000 |
| C3246 | -29.5417140000 | 23.2202240000 | 46.0525620000 |
| C3247 | -29.8388820000 | 23.3599190000 | 47.5127860000 |
| C3248 | -30.1972870000 | 24.1333950000 | 42.6449410000 |
| C3249 | -31.6167540000 | 24.6009970000 | 42.9041570000 |
| C3250 | -32.3965150000 | 24.1039930000 | 43.9641190000 |
| C3251 | -33.7341520000 | 24.4522280000 | 44.0979560000 |
| C3252 | -34.3371750000 | 25.3171480000 | 43.1759450000 |
| C3253 | -36.3333940000 | 26.4155690000 | 42.4577120000 |
| C3254 | -33.5746280000 | 25.8353340000 | 42.1259700000 |
| C3255 | -32.2328190000 | 25.4715700000 | 42.0013840000 |
| C3256 | -28.3407360000 | 25.4079860000 | 42.2277420000 |
| C3257 | -28.0770050000 | 24.8388050000 | 43.4603690000 |
| C3258 | -26.8482470000 | 25.0872240000 | 44.0812330000 |
| C3259 | -25.3732420000 | 24.4902300000 | 45.8603360000 |
| C3260 | -25.9108080000 | 25.8957350000 | 43.4375050000 |
| C3261 | -26.2153500000 | 26.4558100000 | 42.1821030000 |
| C3262 | -25.4455460000 | 27.8542700000 | 40.4037060000 |
| C3263 | -27.4403000000 | 26.2258690000 | 41.5456150000 |
| N3264 | -30.6098540000 | 19.3491680000 | 43.1302930000 |
| N3265 | -29.7407210000 | 24.3220740000 | 45.2222130000 |

| | | | |
|---|---|---|---|
| N3266 | -29.1639720000 | 22.1293790000 | 45.4907620000 |
| O3267 | -28.6361090000 | 20.0468550000 | 42.2865990000 |
| O3268 | -31.7082550000 | 19.6666610000 | 43.9435760000 |
| O3269 | -27.6502020000 | 22.1282410000 | 43.6671570000 |
| O3270 | -35.6597880000 | 25.5842510000 | 43.3903560000 |
| O3271 | -29.5706920000 | 25.0854800000 | 41.7375260000 |
| O3272 | -26.6806310000 | 24.5234000000 | 45.3074210000 |
| O3273 | -25.2223040000 | 27.2350020000 | 41.6606770000 |
| H42 | -34.3468100000 | 21.7674590000 | 40.2274600000 |
| H43 | -32.8345240000 | 22.3282250000 | 42.0852240000 |
| H44 | -31.0257370000 | 21.8387960000 | 37.4982730000 |
| H45 | -33.4578910000 | 21.5193440000 | 37.9162050000 |
| H46 | -29.5094100000 | 22.3900370000 | 39.3728640000 |
| H47 | -27.6993770000 | 26.6405420000 | 40.5818230000 |
| H48 | -24.9450040000 | 26.1204740000 | 43.8698120000 |
| H49 | -34.0088750000 | 26.5077820000 | 41.3959460000 |
| H50 | -31.9634660000 | 23.4361550000 | 44.6953870000 |
| H51 | -34.3347910000 | 24.0640200000 | 44.9140360000 |
| H52 | -31.6570610000 | 25.8639320000 | 41.1722880000 |
| H53 | -31.0081080000 | 17.3277350000 | 42.7206820000 |
| H54 | -29.3031940000 | 17.8351910000 | 42.6211460000 |
| H55 | -30.0858630000 | 17.6380890000 | 44.2192450000 |
| H56 | -33.7178750000 | 19.8156880000 | 43.9001130000 |
| H57 | -32.9538510000 | 19.9434850000 | 42.2828160000 |
| H58 | -33.1044550000 | 18.3501150000 | 43.0886160000 |
| H59 | -30.9925570000 | 21.7978660000 | 43.7222400000 |
| H60 | -29.0732620000 | 22.6960710000 | 41.5494830000 |
| H61 | -29.1947070000 | 24.1260550000 | 47.9602250000 |
| H62 | -30.8780570000 | 23.6698270000 | 47.6681980000 |
| H63 | -29.6630590000 | 22.4085230000 | 48.0151220000 |
| H64 | -36.3121650000 | 25.9901380000 | 41.4453620000 |
| H65 | -37.3681590000 | 26.4768000000 | 42.7994200000 |
| H66 | -35.9040910000 | 27.4258830000 | 42.4278170000 |
| H67 | -25.4459270000 | 23.8824050000 | 46.7634450000 |
| H68 | -24.6616210000 | 24.0275140000 | 45.1662280000 |
| H69 | -25.0183580000 | 25.4947320000 | 46.1267200000 |
| H70 | -24.5411780000 | 28.4243420000 | 40.1838310000 |
| H71 | -25.6078150000 | 27.1136120000 | 39.6095920000 |
| H72 | -26.3047100000 | 28.5374570000 | 40.4353850000 |
| H73 | -29.5343540000 | 25.2423790000 | 45.5875310000 |
| H74 | -27.6753850000 | 21.2953930000 | 43.1561980000 |

Coordinates and gas phase energy (MO6-2X/6-31g++**) for the lowest-energy member of cluster **ADR V**

Gas phase energy...................... -1891.23463161301 hartrees

| atom | x | y | z |
|---|---|---|---|
| C3233 | -29.1871010000 | 20.3319350000 | 42.0119180000 |
| C3234 | -29.4027480000 | 18.5915510000 | 43.8655220000 |
| C3235 | -26.9516470000 | 18.3188040000 | 41.8263100000 |
| C3236 | -29.9985390000 | 21.3343940000 | 42.8328970000 |
| C3237 | -30.7781510000 | 22.3316540000 | 41.9269140000 |

| | | | |
|---|---|---|---|
| C3238 | -32.2575320000 | 21.9978330000 | 41.9625110000 |
| C3239 | -33.0430900000 | 22.2661980000 | 43.0928020000 |
| C3240 | -34.3882260000 | 21.9081200000 | 43.1222290000 |
| C3241 | -34.9713920000 | 21.2686820000 | 42.0258130000 |
| C3242 | -34.1961940000 | 20.9879890000 | 40.9020370000 |
| C3243 | -32.8482050000 | 21.3509430000 | 40.8730600000 |
| C3244 | -29.0584410000 | 22.1506700000 | 43.7855080000 |
| C3245 | -29.6389590000 | 23.6407520000 | 43.7533230000 |
| C3246 | -29.9860350000 | 22.5534530000 | 45.7443660000 |
| C3247 | -30.4387260000 | 22.4502310000 | 47.1653150000 |
| C3248 | -30.3718430000 | 23.7639500000 | 42.3842480000 |
| C3249 | -31.4206320000 | 24.8535430000 | 42.2393970000 |
| C3250 | -32.2177770000 | 24.8929630000 | 41.0901630000 |
| C3251 | -33.1564300000 | 25.8992240000 | 40.8748400000 |
| C3252 | -33.3073380000 | 26.9167520000 | 41.8243080000 |
| C3253 | -35.0716290000 | 27.9802030000 | 40.6120940000 |
| C3254 | -32.4915330000 | 26.9196120000 | 42.9604790000 |
| C3255 | -31.5616710000 | 25.9060570000 | 43.1562440000 |
| C3256 | -28.3716860000 | 24.8922990000 | 42.2689170000 |
| C3257 | -28.5745230000 | 24.6876690000 | 43.6261100000 |
| C3258 | -27.7337660000 | 25.3074520000 | 44.5488310000 |
| C3259 | -27.0793670000 | 25.5561520000 | 46.8362230000 |
| C3260 | -26.7204900000 | 26.1491950000 | 44.0825160000 |
| C3261 | -26.5467640000 | 26.3368390000 | 42.6987470000 |
| C3262 | -25.2801190000 | 27.4407380000 | 40.9985570000 |
| C3263 | -27.3718130000 | 25.7096240000 | 41.7537840000 |
| N3264 | -29.0233470000 | 19.0947000000 | 42.5557480000 |
| N3265 | -30.4442870000 | 23.6200510000 | 44.9688470000 |
| N3266 | -29.2274600000 | 21.6987120000 | 45.1597200000 |
| O3267 | -28.6736580000 | 20.6550780000 | 40.9399010000 |
| O3268 | -28.3759480000 | 18.1375770000 | 41.7686750000 |
| O3269 | -27.7042420000 | 22.0538260000 | 43.4407000000 |
| O3270 | -34.2023490000 | 27.9435210000 | 41.7339540000 |
| O3271 | -29.2443490000 | 24.1728080000 | 41.4977940000 |
| O3272 | -27.9908320000 | 25.0521490000 | 45.8685070000 |
| O3273 | -25.5261630000 | 27.1820970000 | 42.3731710000 |
| H42 | -34.6383060000 | 20.4865150000 | 40.0456320000 |
| H43 | -32.2451680000 | 21.1273070000 | 39.9964600000 |
| H44 | -34.9860110000 | 22.1331110000 | 44.0014350000 |
| H45 | -36.0216370000 | 20.9917410000 | 42.0494050000 |
| H46 | -32.5936170000 | 22.7720840000 | 43.9428950000 |
| H47 | -27.2649090000 | 25.8400910000 | 40.6864690000 |
| H48 | -26.0491480000 | 26.6755260000 | 44.7474210000 |
| H49 | -32.6007830000 | 27.7280690000 | 43.6756760000 |
| H50 | -32.1180270000 | 24.1148400000 | 40.3422670000 |
| H51 | -33.7603760000 | 25.8765060000 | 39.9759210000 |
| H52 | -30.9212930000 | 25.9669750000 | 44.0267090000 |
| H53 | -29.5556780000 | 19.4171850000 | 44.5610800000 |
| H54 | -30.2965180000 | 17.9599790000 | 43.8042260000 |
| H55 | -28.5742780000 | 17.9887510000 | 44.2479950000 |
| H56 | -26.5880910000 | 18.2699330000 | 42.8604620000 |
| H57 | -26.5367720000 | 17.4873670000 | 41.2519270000 |
| H58 | -26.6700620000 | 19.2711040000 | 41.3710840000 |

| | | | |
|---|---|---|---|
| H59 | -30.7205390000 | 20.8176950000 | 43.4638480000 |
| H60 | -30.4110150000 | 22.2241900000 | 40.9039470000 |
| H61 | -30.1202760000 | 23.3368010000 | 47.7260990000 |
| H62 | -31.5313940000 | 22.3972020000 | 47.2197930000 |
| H63 | -30.0077310000 | 21.5616040000 | 47.6265540000 |
| H64 | -35.7053070000 | 27.0848750000 | 40.5605160000 |
| H65 | -35.7034560000 | 28.8595120000 | 40.7490060000 |
| H66 | -34.5163790000 | 28.0794800000 | 39.6699710000 |
| H67 | -26.0603830000 | 25.1956170000 | 46.6507600000 |
| H68 | -27.0757910000 | 26.6536190000 | 46.8557920000 |
| H69 | -27.4254760000 | 25.1805900000 | 47.8007060000 |
| H70 | -24.4352170000 | 28.1307930000 | 40.9691910000 |
| H71 | -25.0175830000 | 26.5240460000 | 40.4547390000 |
| H72 | -26.1467730000 | 27.9088100000 | 40.5136570000 |
| H73 | -30.5539560000 | 24.4994020000 | 45.4578290000 |
| H74 | -27.6389570000 | 22.1485160000 | 42.4769260000 |

Coordinates and gas phase energy (MO6-2X/6-31g++**) for the lowest-energy member of cluster **ADR VI**

Gas phase energy.....................   -1891.24898834473 hartrees

| atom | x | y | z |
|---|---|---|---|
| C1 | -26.4798590000 | 21.5300730000 | 42.6475410000 |
| C2 | -32.6960200000 | 22.8040380000 | 38.9761360000 |
| C3 | -31.7205180000 | 23.0460750000 | 39.9380020000 |
| C4 | -30.4741130000 | 23.9308600000 | 42.5998930000 |
| C5 | -31.8332670000 | 24.5798360000 | 42.7356090000 |
| C6 | -32.9242680000 | 23.8581340000 | 43.2487090000 |
| C7 | -34.1521470000 | 24.4677850000 | 43.4550630000 |
| C8 | -34.3287820000 | 25.8249380000 | 43.1471320000 |
| C9 | -35.8210090000 | 27.6892290000 | 43.0766460000 |
| C10 | -33.2548470000 | 26.5578550000 | 42.6351560000 |
| C11 | -32.0208840000 | 25.9299030000 | 42.4387220000 |
| C12 | -27.8999270000 | 19.0424360000 | 41.5645820000 |
| C13 | -28.4206970000 | 24.9047240000 | 42.2928680000 |
| C14 | -27.3518760000 | 25.5125540000 | 41.6350400000 |
| C15 | -26.1635730000 | 25.6280460000 | 42.3634890000 |
| C16 | -25.0222730000 | 26.6649550000 | 40.5379940000 |
| C17 | -26.0588820000 | 25.1610270000 | 43.6884350000 |
| C18 | -27.1564400000 | 24.5557530000 | 44.3004650000 |
| C19 | -25.9524840000 | 24.1958530000 | 46.3475250000 |
| C20 | -28.3502760000 | 24.4052750000 | 43.5848300000 |
| C21 | -29.6776600000 | 23.8364750000 | 43.9762810000 |
| C22 | -30.9905410000 | 23.5364380000 | 45.8282120000 |
| C23 | -29.6447270000 | 20.1820730000 | 42.8808070000 |
| C24 | -31.9689630000 | 23.9647600000 | 46.8749650000 |
| C25 | -29.7588810000 | 22.3171010000 | 44.4309370000 |
| C26 | -30.3212630000 | 21.5137380000 | 43.2278690000 |
| C27 | -30.4395730000 | 22.4763460000 | 42.0232240000 |
| C28 | -31.5215960000 | 22.1561800000 | 41.0054290000 |
| C29 | -32.3100550000 | 21.0014410000 | 41.0663510000 |
| C30 | -33.2867160000 | 20.7571750000 | 40.0970840000 |

| | | | |
|---|---|---|---|
| C31 | -33.4892620000 | 21.6569210000 | 39.0539270000 |
| N32 | -28.3605740000 | 20.1600470000 | 42.3730760000 |
| N33 | -30.2893400000 | 24.5025950000 | 45.1171270000 |
| N34 | -30.7303380000 | 22.3118840000 | 45.5356410000 |
| O35 | -27.7610400000 | 21.3820720000 | 42.0232630000 |
| O36 | -30.2647950000 | 19.1296050000 | 43.0032060000 |
| O37 | -35.5763390000 | 26.3252560000 | 43.3834980000 |
| O38 | -29.6513990000 | 24.7666280000 | 41.7254490000 |
| O39 | -25.0211200000 | 26.1914350000 | 41.8776060000 |
| O40 | -27.1501350000 | 24.0819090000 | 45.5857520000 |
| O41 | -28.5385400000 | 21.7665730000 | 44.8494580000 |
| H42 | -32.1562810000 | 20.2684320000 | 41.8506370000 |
| H43 | -33.8874730000 | 19.8542370000 | 40.1627370000 |
| H44 | -32.8385440000 | 23.5118340000 | 38.1639170000 |
| H45 | -34.2522900000 | 21.4662710000 | 38.3043580000 |
| H46 | -31.1074770000 | 23.9399590000 | 39.8747870000 |
| H47 | -31.1939540000 | 26.5012090000 | 42.0326280000 |
| H48 | -33.3612500000 | 27.6060140000 | 42.3825690000 |
| H49 | -32.8154690000 | 22.8086650000 | 43.4929060000 |
| H50 | -34.9972610000 | 23.9097580000 | 43.8439410000 |
| H51 | -27.4671200000 | 25.8681460000 | 40.6209940000 |
| H52 | -25.1096790000 | 25.2894180000 | 44.1902220000 |
| H53 | -25.7492750000 | 20.8306960000 | 42.2219040000 |
| H54 | -26.1785500000 | 22.5541590000 | 42.4222170000 |
| H55 | -26.5673410000 | 21.3928080000 | 43.7252900000 |
| H56 | -36.8655450000 | 27.8734660000 | 43.3330700000 |
| H57 | -35.1815840000 | 28.3596300000 | 43.6666680000 |
| H58 | -35.6687970000 | 27.8987620000 | 42.0095380000 |
| H59 | -27.9915970000 | 19.2856690000 | 40.4992470000 |
| H60 | -26.8562050000 | 18.8096400000 | 41.7927850000 |
| H61 | -28.5234850000 | 18.1834840000 | 41.8068200000 |
| H62 | -24.0189730000 | 27.0532600000 | 40.3559890000 |
| H63 | -25.2340150000 | 25.8576380000 | 39.8250000000 |
| H64 | -25.7536820000 | 27.4708300000 | 40.3945880000 |
| H65 | -26.1768160000 | 23.7615040000 | 47.3226770000 |
| H66 | -25.1277110000 | 23.6425180000 | 45.8818250000 |
| H67 | -25.6605570000 | 25.2449140000 | 46.4742250000 |
| H68 | -31.5084820000 | 24.6725380000 | 47.5720270000 |
| H69 | -32.8200940000 | 24.4648700000 | 46.3978700000 |
| H70 | -32.3290040000 | 23.0924030000 | 47.4205950000 |
| H71 | -31.3110100000 | 21.1898370000 | 43.5488550000 |
| H72 | -29.4877720000 | 22.4482020000 | 41.4925290000 |
| H73 | -30.7310550000 | 25.3989280000 | 44.9536180000 |
| H74 | -28.1671250000 | 22.3653160000 | 45.5160450000 |

Coordinates and gas phase energy (MO6-2X/6-31g++**) for the lowest-energy member of cluster **ADR VII**

Gas phase energy...................... -1891.23582482403 hartrees

| atom | x | y | z |
|---|---|---|---|
| C1 | -29.2930430000 | 18.1044180000 | 43.3656430000 |
| C2 | -34.2774370000 | 21.6698100000 | 43.1622290000 |

| | | | |
|---|---|---|---|
| C3 | -32.9872270000 | 22.1811930000 | 43.0387140000 |
| C4 | -30.4333820000 | 23.8252090000 | 42.2722570000 |
| C5 | -31.5151120000 | 24.8944060000 | 42.2198500000 |
| C6 | -31.5217680000 | 25.9842390000 | 43.1063250000 |
| C7 | -32.4658960000 | 26.9968470000 | 43.0078110000 |
| C8 | -33.4369340000 | 26.9581470000 | 42.0007320000 |
| C9 | -35.3514600000 | 27.9903840000 | 41.0130610000 |
| C10 | -33.4245530000 | 25.9062690000 | 41.0793970000 |
| C11 | -32.4668650000 | 24.8991640000 | 41.1958100000 |
| C12 | -27.0090130000 | 19.1551700000 | 41.3519780000 |
| C13 | -28.4049140000 | 24.9144970000 | 42.2214340000 |
| C14 | -27.3662460000 | 25.7149400000 | 41.7780760000 |
| C15 | -26.4725010000 | 26.1844260000 | 42.7520980000 |
| C16 | -24.5089640000 | 27.4913570000 | 43.1761430000 |
| C17 | -26.6053990000 | 25.8521800000 | 44.1118020000 |
| C18 | -27.6668730000 | 25.0215070000 | 44.5079180000 |
| C19 | -26.9416260000 | 24.9738380000 | 46.7942710000 |
| C20 | -28.5873810000 | 24.5914250000 | 43.5657990000 |
| C21 | -29.7114880000 | 23.6106370000 | 43.6447280000 |
| C22 | -30.3553520000 | 22.4483140000 | 45.5233910000 |
| C23 | -28.8289460000 | 20.8008050000 | 41.4772310000 |
| C24 | -30.9757960000 | 22.2871800000 | 46.8749010000 |
| C25 | -29.2058360000 | 22.0879230000 | 43.6605870000 |
| C26 | -29.8654880000 | 21.3970730000 | 42.4304780000 |
| C27 | -30.7996870000 | 22.4198330000 | 41.7208610000 |
| C28 | -32.2406080000 | 21.9613080000 | 41.8733830000 |
| C29 | -32.8195980000 | 21.2135580000 | 40.8411390000 |
| C30 | -34.1107120000 | 20.6985450000 | 40.9641160000 |
| C31 | -34.8454820000 | 20.9255430000 | 42.1270850000 |
| N32 | -28.3035840000 | 19.5762960000 | 41.8613580000 |
| N33 | -30.5574920000 | 23.6346800000 | 44.8309380000 |
| N34 | -29.6800060000 | 21.5320750000 | 44.9266820000 |
| O35 | -28.4511910000 | 19.2476300000 | 43.2266350000 |
| O36 | -28.5273650000 | 21.2988350000 | 40.4032430000 |
| O37 | -34.3343380000 | 27.9873710000 | 42.0032150000 |
| O38 | -29.3363000000 | 24.3380650000 | 41.4149490000 |
| O39 | -25.4745670000 | 26.9836240000 | 42.2712670000 |
| O40 | -27.8664210000 | 24.5750400000 | 45.7923560000 |
| O41 | -27.8081460000 | 21.9722380000 | 43.5631090000 |
| H42 | -32.2495750000 | 21.0322800000 | 39.9330410000 |
| H43 | -34.5414650000 | 20.1226660000 | 40.1496080000 |
| H44 | -34.8440580000 | 21.8580690000 | 44.0703800000 |
| H45 | -35.8520400000 | 20.5289120000 | 42.2257510000 |
| H46 | -32.5616810000 | 22.7761350000 | 43.8398170000 |
| H47 | -32.4822010000 | 24.0954490000 | 40.4696200000 |
| H48 | -34.1496040000 | 25.8547210000 | 40.2762330000 |
| H49 | -30.7586700000 | 26.0696890000 | 43.8707170000 |
| H50 | -32.4694800000 | 27.8325080000 | 43.6999040000 |
| H51 | -27.2318290000 | 25.9713170000 | 40.7356710000 |
| H52 | -25.8969900000 | 26.2222300000 | 44.8372550000 |
| H53 | -29.3593250000 | 17.9308260000 | 44.4419860000 |
| H54 | -30.2955220000 | 18.2893290000 | 42.9613050000 |
| H55 | -28.8635070000 | 17.2228640000 | 42.8721360000 |

| | | | |
|---|---|---|---|
| H56 | -35.9574380000 | 28.8775060000 | 41.2054360000 |
| H57 | -34.9329220000 | 28.0531910000 | 39.9998040000 |
| H58 | -35.9869710000 | 27.0973600000 | 41.0815510000 |
| H59 | -26.9289160000 | 18.0674620000 | 41.4213160000 |
| H60 | -26.9418140000 | 19.4659240000 | 40.3102190000 |
| H61 | -26.2026110000 | 19.6214590000 | 41.9296140000 |
| H62 | -23.8189510000 | 28.0876180000 | 42.5765300000 |
| H63 | -24.9653570000 | 28.1331440000 | 43.9421100000 |
| H64 | -23.9504220000 | 26.6852510000 | 43.6709000000 |
| H65 | -27.2718130000 | 24.4945950000 | 47.7176300000 |
| H66 | -25.9224360000 | 24.6409360000 | 46.5594330000 |
| H67 | -26.9390310000 | 26.0628860000 | 46.9331440000 |
| H68 | -30.7081310000 | 21.3147940000 | 47.2884610000 |
| H69 | -30.6275730000 | 23.0762720000 | 47.5518200000 |
| H70 | -32.0661800000 | 22.3660250000 | 46.8076220000 |
| H71 | -30.4670480000 | 20.5807650000 | 42.8273540000 |
| H72 | -30.5440620000 | 22.4332520000 | 40.6586900000 |
| H73 | -30.5944460000 | 24.4885680000 | 45.3708390000 |
| H74 | -27.4674040000 | 22.0466350000 | 44.4648630000 |

**Supplementary References.**

S1. Ernst, J. T.;  Thompson, P. A.; Nilewski, C.; Sprengeler, P. A.; Sperry, S.; Packard, G.; Michels, T. ; Xiang, A.; Tran, C.; Wegerski, C. J.; Eam, B.; Young, N. P.; Fish, S.; Chen, J.; Howard, H.; Staunton, J.; Molter, J.; Clarine, J.; Nevarez, A.; Chiang, G. G.; Appleman, J. R.; Webster, K. R.; Reich, S. H. *J. Med. Chem*. **2020**, *63*, 11, 5879–5955.
